# QT-GILD: Quartet based gene tree imputation using deep learning improves phylogenomic analyses despite missing data

**DOI:** 10.1101/2021.11.03.467204

**Authors:** Sazan Mahbub, Shashata Sawmya, Arpita Saha, Rezwana Reaz, M. Sohel Rahman, Md. Shamsuzzoha Bayzid

**Affiliations:** Department of Computer Science and Engineering Bangladesh University of Engineering and Technology, Dhaka-1205, Bangladesh; Department of Computer Science University of Maryland, College Park, Maryland 20742, USA

**Author notes:** These authors contributed equally to this work (Sazan Mahbub, Shashata Sawmya).

## Abstract

Species tree estimation is frequently based on phylogenomic approaches that use multiple genes from throughout the genome. However, for a combination of reasons (ranging from sampling biases to more biological causes, as in gene birth and loss), gene trees are often incomplete, meaning that not all species of interest have a common set of genes. Incomplete gene trees can potentially impact the accuracy of phylogenomic inference. We, for the first time, introduce the problem of imputing the quartet distribution induced by a set of incomplete gene trees, which involves adding the missing quartets back to the quartet distribution. We present QT-GILD, an automated and specially tailored unsupervised deep learning technique, accompanied by cues from natural language processing (NLP), which learns the quartet distribution in a given set of incomplete gene trees and generates a complete set of quartets accordingly. QT-GILD is a general-purpose technique needing no explicit modeling of the subject system or reasons for missing data or gene tree heterogeneity. Experimental studies on a collection of simulated and empirical data sets suggest that QT-GILD can effectively impute the quartet distribution, which results in a dramatic improvement in the species tree accuracy. Remarkably, QT-GILD not only imputes the missing quartets but it can also account for gene tree estimation error. Therefore, QT-GILD advances the state-of-the-art in species tree estimation from gene trees in the face of missing data. QT-GILD is freely available in open source form at https://github.com/pythonLoader/QT-GILD.

## 1 Introduction

High-throughput DNA sequencing is generating new genome-wide data sets for phylogenetic analyses, potentially including hundreds or even thousands of loci. The estimation of species trees from multiple genes is necessary since true gene trees can differ from each other and from the true species tree due to various processes, including gene duplication and loss, horizontal gene transfer, and incomplete lineage sorting (ILS) [1]. ILS (also known as deep coalescence) is considered to be a dominant cause for gene tree heterogeneity, which is best understood under the coalescent model [2–9]. In the presence of gene tree heterogeneity, standard methods for estimating species trees, such as concatenation (which combines sequence alignments from different loci into a single “supermatrix”, and then computes a tree on the supermatrix) can be statistically inconsistent [10,11], and produce incorrect trees with high support [12]. Therefore, *summary methods* that operate by combining estimated gene trees and can explicitly take gene tree discordance into account are gaining increasing attention from the systematists, and some of the coalescent based summary methods are statistically consistent [13–22]. Other coalescent speciestree estimation methods include BEST [23] and *BEAST [24], which co-estimate gene trees and species tree from input sequence alignments. These co-estimation methods can produce substantially more accurate trees than other methods, but are computationally intensive and do not scale up for genome-level analyses [25–28].

There has been notable progress in constructing large scale phylogenetic trees by analyzing genomewide data, but substantial challenges remain. Assembling a complete dataset with hundreds of orthologous genes from a large set of species remains a difficult task which has downstream impact on species tree inference [29–31]. Gene trees can be incomplete due to various reasons ranging from biological processes as in gene birth and loss [32] to biases in taxon and gene sampling, stochasticity inherent in collecting data across thousands of loci, and difficulty in sequencing and assembling the complete set of taxa of interest (see [29, 30, 33, 34] for more elaborate discussion). Incomplete gene trees may decrease the species tree accuracy [27, 31, 35] and introduce ambiguity in the tree search [36, 37]. Indeed, as we will show in our experimental results, incomplete gene trees substantially reduce the accuracy of the best existing coalescent based summary methods.

We address the problem of missing data in phylogenomic data sets by formulating the *Quartet Distribution Imputation* (QDI) problem, where we seek to add the missing quartets to the quartet distribution induced by a given set of incomplete gene trees. Quartet-based methods have gained substantial interest as quartets (4-leaf unrooted gene trees) do not contain the “anomaly zone” [2, 38, 39], a condition where the most probable gene tree topology may not be identical to the species tree topology. ASTRAL [14], which is one of the most accurate and widely used coalescent-based methods, seeks to infer a species tree so that the number of induced quartets in the gene trees that are consistent with the species tree is maximized. Another approach is to infer individual quartets (with or without weights), and then amalgamate them into a single coherent species tree [18, 40–46]. wQFM [46] and wQMC [45] represent the latter category of quartet-based methods.

Existing methods for adding missing taxa to the gene trees use a “reference tree” and attempt to add the missing species in such a way that various distance metrics (e.g., “extra-lineage score”, Robinson-Foulds distance) are optimized with respect to the given reference tree [27, 47]. However, obtaining a reasonably accurate reference tree in the presence of missing data is difficult [47], which subsequently affects the tree completion steps as the reference tree can be far from the true tree. In this study, we present QT-GILD (**Q**uar**t**et based **G**ene tree **I**mputation using **D**eep **L**earning) – a novel deep learning based technique which learns the quartet distribution induced by the given set of incomplete gene trees using an especially tailored autoencoder model, and subsequently generates a complete set of quartets. In doing so, it not only imputes the missing quartets, but also attempts to correct the quartets present in the given set of estimated and incomplete gene trees by leveraging the underlying quartet distribution. Therefore, the complete set of quartets generated by QT-GILD is often closer to the set of quartets present in the true gene trees than to those present in the estimated gene trees. Thus, QT-GILD obviates the need for a reference tree as well as accounts for gene tree estimation error. Our experimental results, on a collection of simulated and real biological data sets covering a wide range of model conditions, indeed show that amalgamating the imputed set of quartets generated by QT-GILD can remarkably improve the accuracy of species tree inference. Additionally, we particularly aimed for making QT-GILD an unsupervised machine learning technique. Unsupervised learning approaches have advantages over supervised methods particularly when the data are heterogeneous, which are often so with various phylogenomic datasets and therefore the supervised models trained on a particular set of incomplete gene trees may not be generalizable to a new set of input gene trees. Furthermore, QT-GILD is a general-purpose approach which requires no explicit modeling of the reasons of gene tree heterogeneity or missing data, making it less vulnerable to model mis-specification. QT-GILD is the first of its kind which advances the state-of-the-art in species tree estimation in the presence of missing data.

## 2 Quartet Imputation Problem

### 2.1 Problem Definition

Let 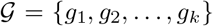 be a set of *k* gene trees, where each *g_i_* is a tree on taxon set *S_i_* ⊆ *S* (i.e., any gene tree *g_i_* can be on the full set *S* of *n* taxa or can be on a subset *S_i_* of taxa, making the gene tree incomplete). For a set of four taxa *a,b,c,d* ∈ *S*, the quartet tree *ab*|*cd* denotes the unrooted quartet tree in which the pair *a, b* is separated from the pair *c, d* by an edge. Let 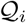 be the set of quartets in *g_i_*. Therefore, 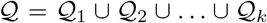 is the multi-set of quartets present in 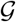. Note that there 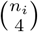 are quartets in 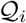, where *n_i_* = |*S_i_*|. When all the trees in 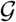 are complete, 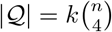. Let 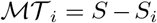 be the set of missing taxa in *g_i_*. Therefore, any quartet *q* involving any subset of taxa in 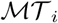 is a missing quartet in *g_i_*. In the presence of missing taxa in the gene trees, 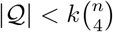. We now define the quartet imputation problem as follows.

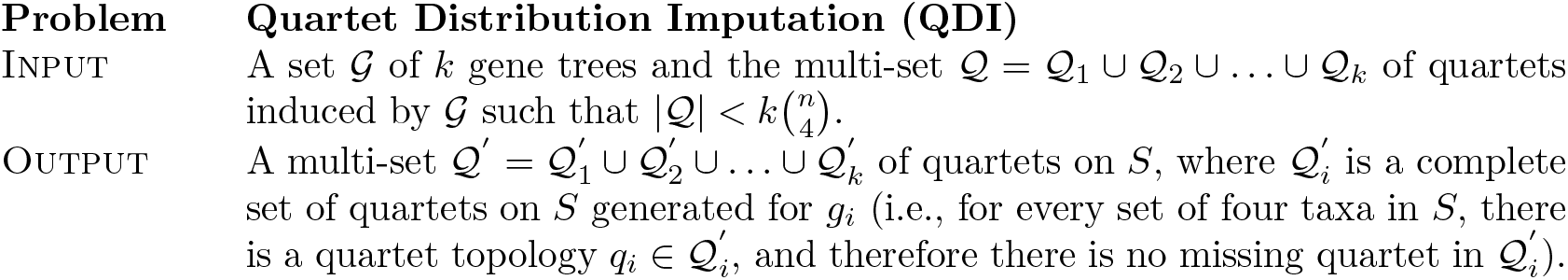

A natural optimization criteria to solve QDI is to impute the missing quartets in 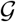 in such a way that the total number of consistent quartets in 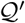 is maximized (a set of quartets is consistent if they can co-exist in a single tree). In this study, we do not approach QDI as a discrete optimization problem; instead, we propose a machine learning approach to learn the underlying quartet distribution in 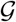 and infer the missing quartets accordingly.

### 2.2 Approach

We discuss various components of QT-GILD (in Sections 2.2.1–2.2.4) followed by the end-to-end pipeline in Section 2.2.5.

#### 2.2.1 Quartet encoding

For the set *S* of *n* taxa present in 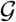, there are 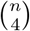 subsets of four different taxa. For a set *qs_j_* 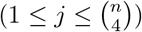 of four taxa *a, b,c,d* ∈ *S*, there are three possible quartets: *ab*|*cd, ac*|*bd*, and *ad*|*bc*. For a set of four taxa *qs_j_* and a gene tree 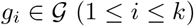, we define a *quartet vector q_i,j_* which represents a probability distribution of the presence of the three possible quartets in *g_i_*. If 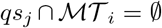 (meaning that *a, b, c, d* are present in *g_i_*), and if 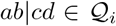 then *q_i,j_* = (1, 0, 0), where 1 represents the presence of *ab*|*cd* and the other two 0s represent the absence of *ac*|*bd* and *ad*|*bc* in 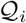. When 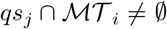 (i.e., there is no quartet on *qs_j_* in 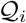), we set 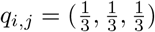, representing a missing quartet which needs to be imputed and equal probabilities of the three possible quartets topologies for imputation. Next, for a particular set *qs_j_* of four taxa, we define a *k* × 3 *quartet encoding* matrix 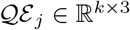 where the *i*-th row represents the quartet vector *q_i,j_*. See Fig. S1 in Supplementary Material for an example.

#### 2.2.2 Masking for self-supervision

We particularly aimed for developing an unsupervised autoencoder based model, but in order to increase the learning capability of our model in a supervised manner, we make it a *self-supervised* model where a portion of the input is used as a supervisory signal to the autoencoder fed with the remaining portion of the input. In this context, we leverage a technique called “masking”, which has been used in several state-of-the-art language modeling approaches, where we mask out a certain percentage of non-missing quartets in a quartet encoding matrix 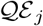, and ask the autoencoder to predict the masked quartets.

We mask out randomly selected *λ*|*S*| (0 ≤ *λ* < 1) non-missing quartets by replacing the one-hot encoded quartet vectors corresponding to the randomly selected non-missing quartets by 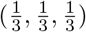. In this study, we set *λ* = 0.1. Thus, we create a masked quartet encoding matrix 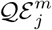 from the original matrix 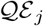. During the training, the autoencoder takes 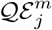 as input and tries to predict the masked quartet vectors. The corresponding cross entropy loss *J* as the reconstruction loss (error) of predicting the masked entries with respect to the original quartet encoding matrix 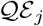 is computed and used as a feedback. Thus, the autoencoder is trained iteratively by backpropagating the loss *J* within a feedback-loop, transforming a fully unsupervised approach into a self-supervised autoencoder and thereby helping the model to effectively learn the quartet distribution.

#### 2.2.3 Positional encoding

Masked quartet encoding matrices 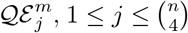 defined on 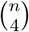 subsets of four taxa 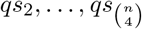 encode the information regarding three possible quartets on different four taxa set across all the gene trees in 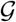. However, 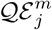 does not encode any information about *qs_j_*, making it impossible for the autoencoder to identify which set of four taxa is represented by a particular 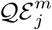. In order to inject some information about different sets of four taxa, we incorporate an extra non-learnable signal, *positional-encoding* (PE) [48]. PE was originally proposed for NLP by Vaswani *et al*. [48] and is widely used to encode positional information of the tokens in sequence data. Recently, it has also been successfully applied to computational biology [49]. For the *j*-th four-taxa-set *qs_j_*, we generate a *k* × 3 positional-encoding matrix 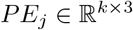 according to Equation [1].

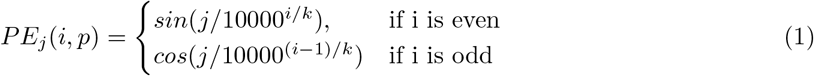

Here, *PE_j_*(*i, p*) represents the positional-encoding of the *p*-th possible quartet (1 ≤ *p* ≤ 3) on *qs_j_* for the *i*-th gene tree 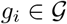 and 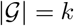. Note that this function is constant with respect to *p* so that we get the same positional-encoding for three possible quartets on *qs_j_* for a particular gene tree *g_i_*. The output of positional encoding *PE_j_* is element-wise added with 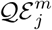 resulting in new representations 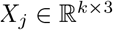 which contain not only the information about three possible quartets across all the gene trees, but also the information about different sets of four-taxa *qs_j_*, 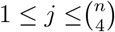 (see Eqn. 2 and Fig. 1 (d)).

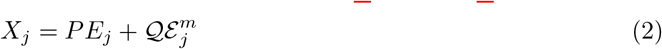

**Figure 1:**
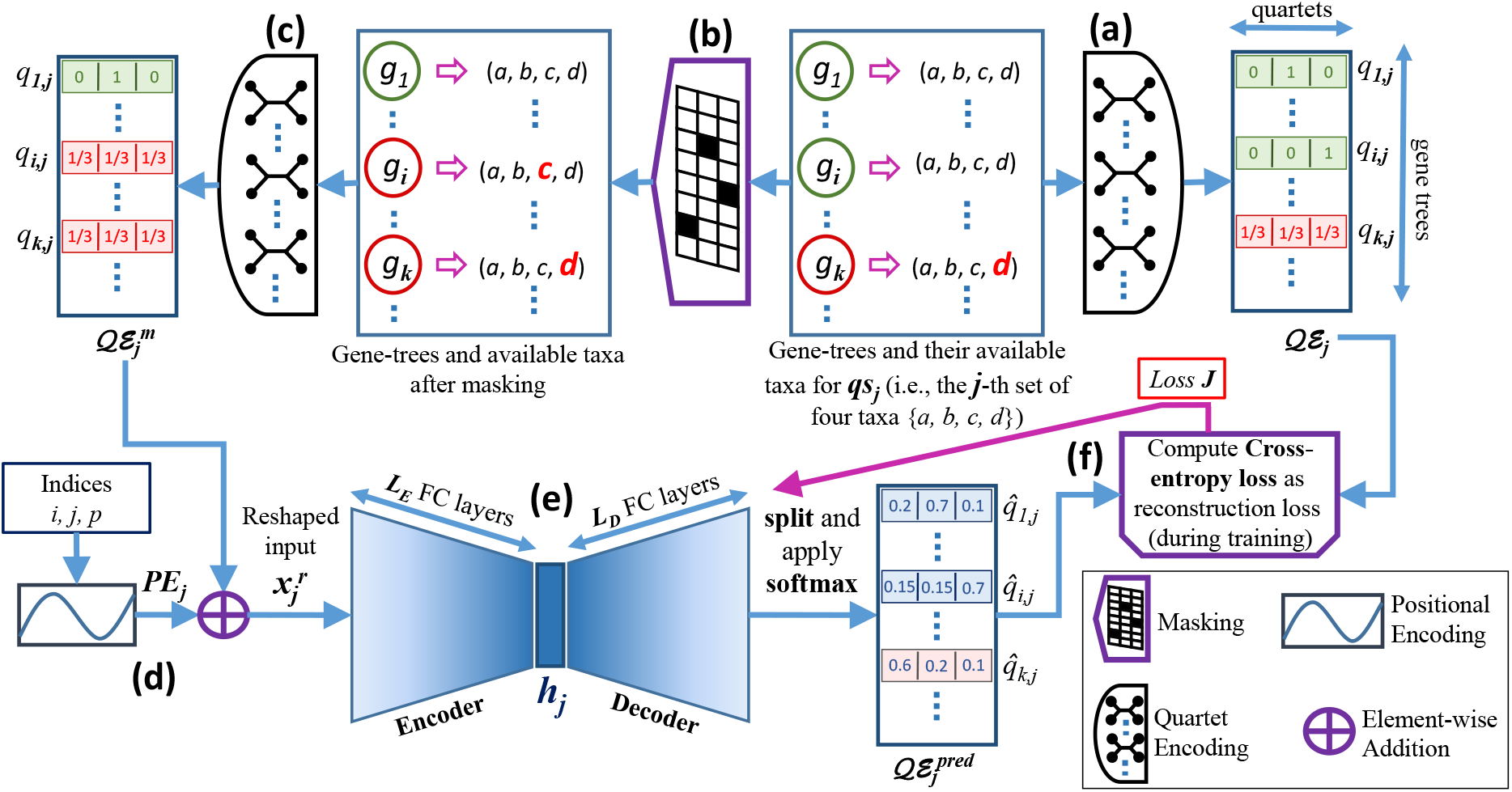
Overall pipeline of QT-GILD. (a) Generation of *Quartet Encoding* matrix 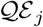 for a set of four taxa, *qs_j_*. (b) Masking (in this example, taxon *c* was masked out from gene tree *g_i_*). (c) Generation of *masked quartet encoding* matrix 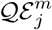 from the output of masking. (d) Positional encoding *PE_j_* generation from indices *i, j, p*, which are the index of a gene tree, index of a set of four taxa, and index of a possible quartet topology, respectively. (e) The reshaped version of the element-wise summation of 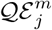 and *PE_j_* (i.e., 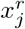) is passed through the autoencoder and the output is split and normalized using softmax, generating 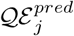. (f) Cross-entropy loss *J* is computed between 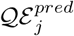 and 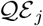 and optimized to update the parameters of the autoencoder (the backward arrow represents backpropagation through the network).

#### 2.2.4 Deep autoencoder

Autoencoder (AE) [50] is a type of artificial neural network that learns to copy its input to its output. The deep autoencoder architecture consists of two modules – encoder and decoder – stacked one after the other and built using fully-connected layers (Figure 1 (e)). The output *X_j_* of the element-wise summation of 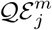 and *PE_j_* (as discussed in Section 2.2.3) is reshaped into a vector 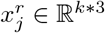, which is used as the input of the encoder module. By using subsequent *L_E_* fully-connected layers, the dimension of 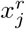 is reduced and a condensed latent representation *h_j_* is generated, which is subsequently used as the input of the decoder module. In the decoder, the dimension of *h_j_* is expanded by *L_D_* fully-connected layers (here, *L_E_* and *L_D_* are two hyperparameters). The output of the *L_D_*-th layer is divided into *k* equal segments of length 3 and softmax normalization is applied on each of them. Here, softmax on the *i*-th segment generates the predicted quartet-vector 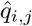, which is the estimated probability distribution over the possible quartets on *qs_j_* for the gene-tree *g_i_*. Thus, 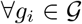, we get 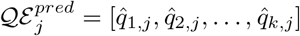, which is the predicted quartet-encoding matrix corresponding to *qs_j_*.

#### 2.2.5 Overall pipeline of QT-GILD

Figure 1 shows the overall end-to-end pipeline of QT-GILD comprising the individual components described in Sections 2.2-2.5. For each set of four taxa *qs_j_*, we first create a quartet encoding matrix 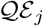, which is subsequently masked to produce 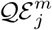. Next, these masked encoding matrices are added to positional encoding matrices and reshaped to generate the input 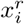 of the autoencoder. The autoencoder network produces the predicted quartet-encoding matrix 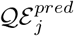. During the training phase, considering 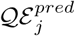 as the prediction and 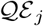 as the ground truth, we compute the crossentropy loss *J* as the reconstruction loss for the non-missing quartets, i.e., for the quartet-vectors 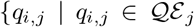 and 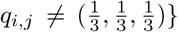. The autoencoder is trained by backpropagating this loss in a self-supervised fashion (see Fig. 1(f) and Section 2.2.2). This training with backpropagation is run for a predefined number of epochs. Finally, it is run once without the loss function to produce the final encoding matrix 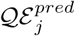. Next, we generate the set of imputed quartets 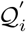 – representing the quartets in the *i*-th gene-tree *g_i_* – where the *j*-th quartet 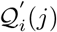 is computed according to Equation [3]. This imputed set of quartets is subsequently used by quartet amalgamation techniques, such as wQFM and wQMC.

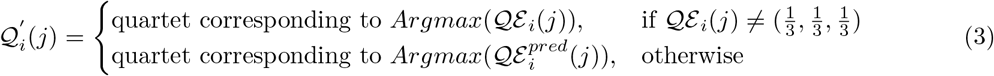

## 3 Experimental Study

### 3.1 Datasets

We used previously studied simulated and biological datasets to evaluate the performance of QT-GILD. We studied three collections of simulated datasets: one based on a biological dataset (37-taxon mammalian dataset) and a 15-taxon dataset that were generated in some prior studies [51,52]. These datasets consist of gene sequence alignments generated under a multi-stage simulation process that begins with a species tree, simulates gene trees down the species tree under the multi-species coalescent model (and so can differ topologically from the species tree), and then simulates gene sequence alignments down the gene trees. These datasets vary from moderately low to extremely high levels of ILS, and also vary in terms of number of genes and gene-tree estimation errors (controlled by sequence lengths). Thus, the simulated datasets provide a range of conditions in which we explored the performance of QT-GILD and investigate the impact of quartet distribution imputation in species tree inference. Table S1 in Supplementary Material presents a summary of these datasets. We also evaluated QT-GILD on a challenging biological dataset comprising 42 angiosperms from Xi *et al*. [53].

### 3.2 Generating incomplete gene trees

We deleted taxa randomly, varying the number of taxa removed from each gene tree, thus producing incomplete gene trees. Instead of a fixed number of taxa, we removed different ranges of taxa. For a particular gene tree *g_i_* in the input set 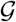 of gene trees and for a particular range *x–y* of missing taxa, we randomly select an integer *mt* ∈ [*x, y*], and randomly select and delete *mt* taxa from *g_i_*. For example, for a range of 3–4 missing taxa, we remove 3 or 4 (selected randomly) from the gene trees in 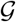. We varied the number of missing taxa (2-40%) for different datasets – creating model conditions with 13%-80% missing quartets. In all the datasets analyzed in this study, this random taxa deletion protocol did not remove any particular taxa from all the gene trees, i.e., each taxa remained present in at least one incomplete gene tree.

### 3.3 Species tree estimation methods

We used wQFM [46], a highly accurate weighted quartet amalgamation technique, to estimate species trees from weighted quartets. We also used wQMC, which is another well known weighted quartet amalgamation technique, in order to show the usability of the imputed quartets generated by QT-GILD across various quartet amalgamation techniques as well as to show that the improvements in species tree inference resulting from amalgamating imputed quartets is not due to wQFM, rather mostly due to the effective imputation of the quartet distribution by QT-GILD. We ran wQFM and wQMC using the embedded quartets in the gene trees with weights reflecting the frequencies of the quartets (i.e., number of gene trees that induce a particular quartet). We compared wQFM with ASTRAL-III [14, 54] (version 5.7.3), which is one of the most accurate and widely used quartet amalgamation techniques. These methods were evaluated on both complete and incomplete gene trees, showing the impact of missing data. wQFM and and wQMC were also run on imputed set of quartets, generated by QT-GILD, to demonstrate the impact of quartet imputation on species tree estimation. Therefore, we evaluated the following variants of different methods.

- *ASTRAL-complete*: ASTRAL, when run on a given set of complete gene trees.
- *ASTRAL-incomplete*: ASTRAL, run on a given set of incomplete gene trees.
- *wQFM-complete*: wQFM, run on the set of weighted quartets induced by a given set of complete gene trees.
- *wQFM-incomplete*: wQFM, run on the set of weighted quartets induced by a given set of incomplete gene trees.
- *wQFM-imputed*: wQFM, run on the imputed set of weighted quartets generated by QT-GILD from a given set of incomplete gene trees.
- wQMC-complete, wQMC-incomplete and wQMC-imputed: defined similarly as wQFM-complete, wQFM-incomplete and wQFM-imputed.

Note that ASTRAL cannot take a set of quartets as input and as such we cannot evaluate ASTRAL on imputed quartet distributions. For brevity, we denote by *complete quartet distributions* and *incomplete quartet distributions* the weighted quartet distributions induced by complete and incomplete estimated gene trees, respectively. Because wQFM generally produces better trees than wQMC [46] and to keep the figures and relevant discussion readable and easy to follow, the results for ASTRAL and wQFM are presented here, while the results for wQMC are presented in the Supplementary Material.

### 3.4 Measurements

We compared the estimated trees (on simulated datasets) with the model species tree using normalized Robinson-Foulds (RF) distance [55] to measure the tree error. The RF distance between two trees is the sum of the bipartitions (splits) induced by one tree but not by the other, and vice versa. We also compared the quartet scores (the number of quartets in the gene trees that agree with a species tree) of the trees estimated by different methods. All the trees estimated in this study are binary and so False Positive (FP), and False Negative (FN) and RF rates are identical. For the biological dataset, we compared the estimated species trees to the scientific literature. We analyzed multiple replicates of data for various model conditions and performed two-sided Wilcoxon signed-rank test (with *α* = 0.05) to measure the statistical significance of the differences between two methods. We assessed the quality of the quartet distributions, induced by complete, incomplete, and imputed gene tree distributions, by comparing them with the true quartet distribution (i.e., the quartets induced by true gene trees).

## 4 Results and Discussion

### 4.1 Results on 15-taxon dataset

The average RF rates of different variants of ASTRAL and wQFM on various model conditions in 15-taxon dataset are shown in Fig. 2. We have investigated the performance on varying gene tree estimation errors using 100bp and 1000bp sequence lengths and on varying numbers of genes (100 and 1000). In general, wQFM is more accurate than ASTRAL both on complete and incomplete gene trees. For all levels of missing data, certain trends were clearly seen. As expected, tree accuracy of ASTRAL and wQFM deteriorated in the presence of missing taxa, and the RF rates increased with increasing levels of missing data. However, tree accuracy of ASTRAL is more impacted by increasing levels of missing data than that of wQFM as wQFM-incomplete was notably better than ASTRAL-incomplete on most of the model conditions (especially, see the results on 100gene-1000bp and 1000gene-100bp model conditions) and in many cases, the differences are statistically significant (*p* ≪ 0.05). ASTRAL and wQFM improves in accuracy as the number of genes increased (from 100 to 1000) and often achieved competitive tree accuracy (compared to the accuracy of the trees estimated on complete gene trees) even when the amount of missing data was large. This is mostly due to the fact that a large number of gene trees provide more unique bipartitions (and hence more quartets) than a relatively smaller number of genes. Similar trends were observed in case of wQMC (see Sec. 5 in Supplementary Material). wQMC is, in general, less accurate than wQFM both on complete and incomplete gene trees. Notably, WQMC-incomplete is slightly better than ASTRAL-incomplete in some model conditions (e.g., 100gene-100bp and 1000gene-100bp model conditions with 4–5 and 5–6 missing taxa in Supplementary Fig. S3).

**Figure 2:**
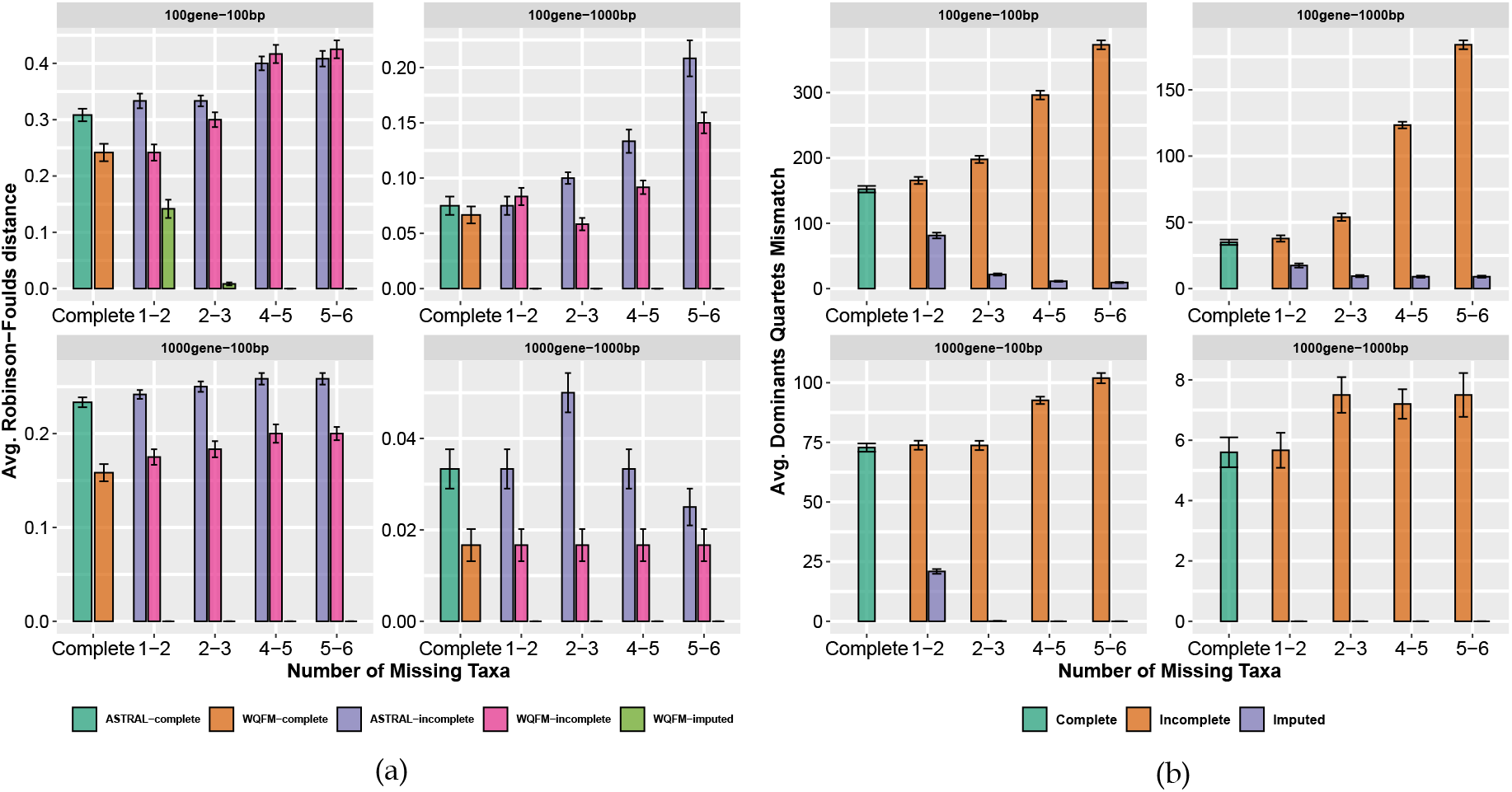
Results on 15-taxon dataset. (a) Comparison of different variants of ASTRAL and wQFM on 15-taxon dataset. We show the average RF rates with standard errors over 10 replicates. For each model condition, we varied the taxa removal rate from ~6% to ~40%, resulting in 13-80% missing quartets. (b) Average numbers of dominant quartets (out of 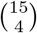) in different quartets distributions that differ from the true dominant quartets.

The most important and interesting results were observed on imputed sets of quartets. wQFM-imputed substantially outperformed not only wQFM-incomplete and ASTRAL-incomplete but also wQFM-complete and ASTRAL-complete, showing the efficacy of QT-GILD in imputing quartet distributions. The improvements are remarkable as in most of the model conditions, wQFM-imputed returned the true species tree (RF-rate = 0), whereas ASTRAL-incomplete and wQFM-incomplete incurred as high as ~45% errors (note the higher taxa removal rates on 100gene-100bp model condition). This dramatic improvement was also observed in case of wQMC-imputed. Even though wQMC-imputed is not as good as wQFM-imputed trees in some cases (see 100gene and 1000gene-1000bp model conditions with smaller numbers of missing taxa in Fig. S3 in the Supplementary Material), wQMC-imputed trees returned the true species trees in most of the cases and consistently outperformed ASTRAL-incomplete as well as ASTRAL-complete. Note that, even with a reasonably well imputed set of quartets, the expected trend is for the accuracy of the tree estimated on an imputed set of quartets to be higher than that of the tree estimated on the corresponding incomplete quartet distribution but lower than or as good as the tree accuracy obtained on the complete quartet distribution. But, surprisingly, wQFM-imputed and wQMC-imputed improves upon wQFM-complete and wQMC-complete (as well as ASTRAL-complete) on this particular dataset. These results suggest, and as we will show in the following, that the imputed quartet distributions are closer to true quartet distributions (i.e., the quartet distribution induced by true gene trees) than the complete and incomplete quartet distributions are to true distributions. Note that, on the 100gene-100bp model condition, the RF rates of the trees estimated on imputed quartets with 1-2 missing taxa were higher than those with larger amounts of missing data. We believe this is because there are only 18272.8 missing quartets with 1-2 missing taxa, and thus QT-GILD could not account for gene tree estimation error (while imputing the missing quartets) as much as it did in other model conditions with larger numbers of missing quartets.

We performed a series of experiments to show the impact and quality of the quartet distributions produced by QT-GILD. First, we measure the divergence between true quartet distributions and different sets of quartet distributions in estimated gene trees (e.g., complete, incomplete and imputed) in terms of the number of “dominant” quartets that differ between two quartet distributions. For a set of four taxa *a, b, c, d*, the dominant quartet (out of three possible quartets *ab*|*cd*, *ac*|*bd*, and *bc*|*ad*) is defined to be the quartet with the highest weight (i.e., the most frequent quartet topology). A dominant quartet topology is the statistically consistent estimate for the true evolutionary history among a group of four taxa since there are no *anomalous* 4-leaf unrooted gene trees [39, 56]. For a set of *n* taxa, there are 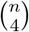 different four-taxa sets, and thus 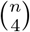 dominant quartets. In Fig. 2(b), we show the numbers of dominant quartets (out of 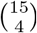) in complete, incomplete and imputed set of quartets that differ from the dominant quartets in true gene trees. Incomplete quartet distributions had the highest numbers of mismatches, followed by complete quartet distribution and the imputed sets of quartets incurred the lowest amount of mismatch (in most cases, the numbers of mismatches are ~0), explaining why the RF rates of the trees estimated from imputed quartets are ~0. The numbers of mismatches in incomplete quartet distributions increased as we increase the amount of missing data. In contrast, the numbers of mismatch in imputed quartet distributions decreased with increasing amounts of missing data (especially on model conditions with higher amounts of gene tree estimation error, ie., the 100-bp model condition). This is because, in the presence of higher numbers of missing quartets, QT-GILD had the opportunity to impute more quartets and in doing so it accounted for larger amounts of gene tree estimation error. This explains the seemingly counter-intuitive trend that RF rates of wQFM-imputed decreased with increasing amounts of missing data. Next, we measured the divergence between the quartet distribution of estimated gene trees and the quartet distribution of true gene trees using Jensen-Shannon divergence [57]. We represent the gene tree distribution by the frequency of each of the three possible alternative topologies for all the 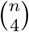 quartets of taxa. Jensen-Shannon divergence of complete, incomplete and imputed quartet distributions from true quartet distributions are shown in Fig. S2 in Supplementary Material. The fact that the difference (both in terms of numbers of mismatch in dominant quartet topologies and Jensen-Shannon distance) between true and complete quartet distributions is higher than the difference between true and imputed quartet distributions suggests that QT-GILD not only imputes the missing quartets, but also accounts for gene tree estimation error. Next, in order to show the efficacy of QT-GILD in imputing missing quartets, we report the proportion of missing quartets that were correctly imputed, with respect to both estimated and true gene trees, by QT-GILD (see Table S2 in Supplementary Material). QT-GILD was able to correctly impute around 51-58% and 51-63% of missing quartets with respect to estimated and true gene trees, respectively.

Finally, we computed the quartet scores of different species trees estimated by different methods with respect to both estimated and true gene trees (see Supplementary Tables S3 and S4.). Interestingly, on estimated gene trees, although ASTRAL obtained higher quartet scores than wQFM-incomplete, and wQFM-imputed in most of the model conditions, the scores of wQFM-incomplete and wQFM-imputed are closer to the true quartet score than the scores of ASTRAL-estimated trees are to the true score. This is mostly a result of the presence of gene tree estimation error. The statistical consistency of estimating species trees by maximizing the quartet score criterion holds when we have a sufficiently large number of true gene trees (with no estimation error). Unfortunately, in practice, the number of genes is limited and the estimated gene trees are not error-free. Therefore, the optimal tree with respect to the quartet score may not be the true species tree. As such, quartet-based methods may “overshoot” the quartet score as they return trees with higher quartet scores than the true quartet score, especially when we have a limited number of estimated gene trees (with estimation error) [37]. The improved performance of wQFM and the efficacy of QT-GILD as an imputation method are even more evident from the quartet scores when they are computed with respect to the true gene trees (i.e., no estimation error). Across all the model conditions, wQFM-imputed obtained the highest quartet scores (w.r.t true gene trees), followed by wQFM-complete and ASTRAL-complete (Table S4 in Supplementary Material). The lowest scores were obtained when the gene trees are incomplete. Notably, wQFM-incomplete consistently achieved higher quartet scores than ASTRAL-incomplete. Note that the quartet consistency score is a statistical consistent measure meaning that the higher the quartet scores, the more accurate the species trees will be (given a sufficiently large number of true gene trees). Therefore, these quartet scores with respect to true gene trees support the trends observed in RF rates (see Fig. 2).

### 4.2 Results on 37-taxon mammalian simulated dataset

In the simulated mammalian data, we explored the impact of varying numbers of genes (25 - 800), varying amounts of gene tree estimation error (i.e., the amount of phylogenetic signal by varying the sequence length for the markers: 250bp - 1500bp). We also investigated three levels of ILS (shorter branches increases ILS) by multiplying or dividing all internal branch lengths in the model species tree by two – producing three model conditions that are referred to as 1X (moderate ILS), 0.5X (high ILS) and 2X (low ILS). For each model condition, we varied the taxa removal rate from ~2% to ~10%, resulting in 6 - 25% missing quartets. Because the number of quartets grows exponentially with the number of taxa, even with 25% missing quartets, the 37-taxon dataset contained a substantial number (~7,00,000 - ~33,00,000) of missing quartets.

Fig. 3(a) shows the RF rates of various methods on varying ILS levels (0.5X, 1X, 2X) with 200 genes and a fixed sequence length (500 bp). As expected, species tree error rates of various methods increased as ILS level increased. Similar to 15-taxon dataset, wQFM is better than ASTRAL (both on complete and incomplete datasets), and wQFM-imputed is consistently better than wQFM-incomplete. Remarkably, for moderate (1X) to low ILS (2X) datasets, wQFM-imputed is substantially better than ASTRAL-complete and wQFM-complete (except for 3-4 taxa removal rate on 2X model condition). On the high-ILS dataset (0.5X), wQFM-imputed is better than wQFM-incomplete and ASTRAL-incomplete. Moreover, for small amount of missing taxa (1-2), wQFM-imputed is even better than ASTRAL-complete and wQFM-complete. In some cases with small numbers of missing taxa, wQFM-incomplete is better than wQFM-complete (albeit the differences are small and not statistically significant). The improved performance resulting from the imputed set of quartets was also observed for wQMC. wQMC-imputed was as good as or better than wQMC-incomplete (see Fig. S4 in the Supplementary Material).

**Figure 3:**
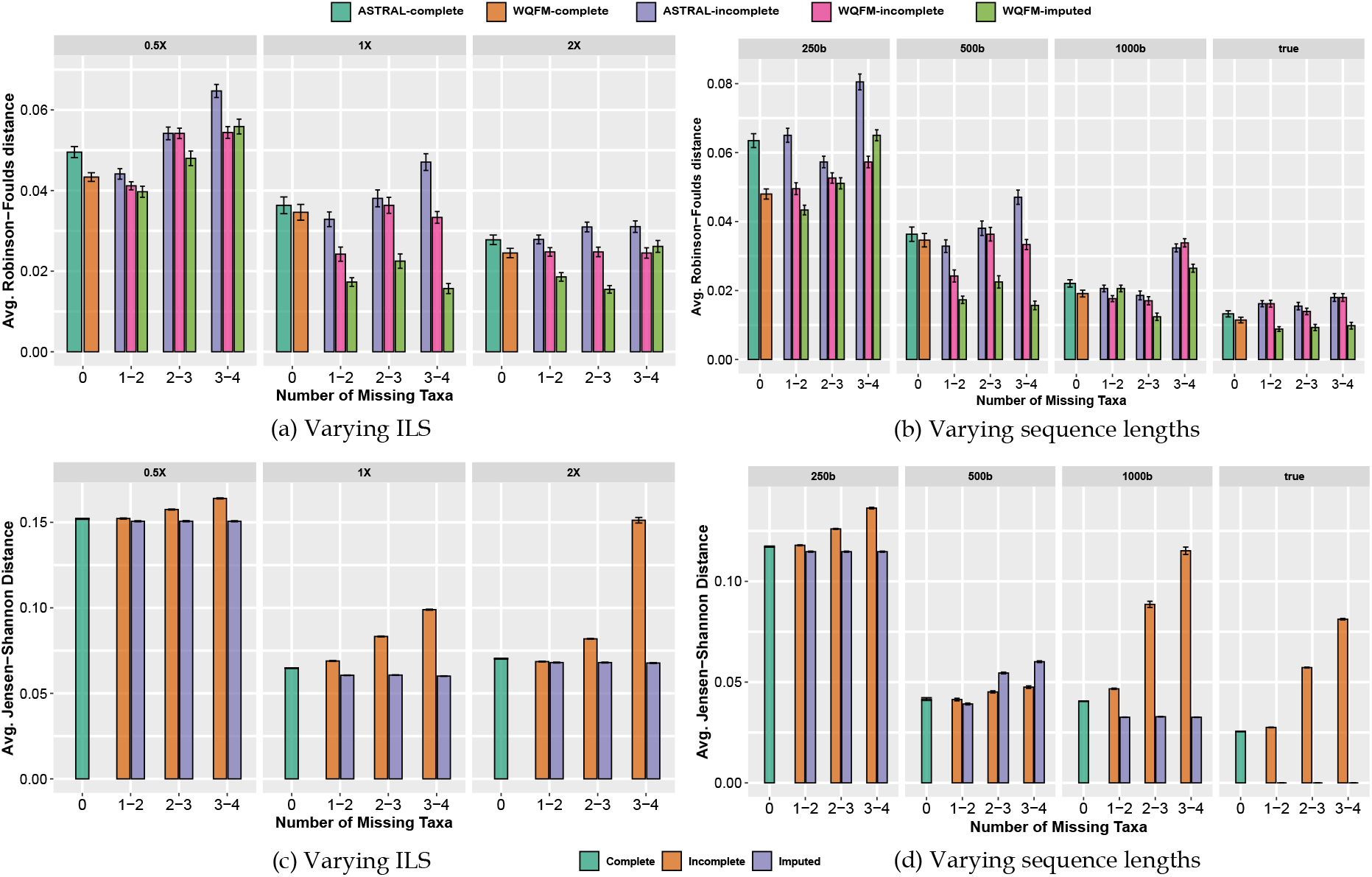
Results on 37-taxon dataset. (a)-(b) We show the average RF rates with standard errors over 20 replicates. (a) The level of ILS was varied from 0.5X (highest) to 2X (lowest) amount, keeping the sequence length fixed at 500bp and the number of genes at 200. (b) The sequence length was varied from 250bp to 1500bp, keeping the number of genes fixed at 200, and ILS at 1X (moderate ILS). We also analyzed the true gene trees. (c)-(d) Jansen-Shannon divergence between true quartet distributions and complete, incomplete and imputed quartet distributions.

RF rates on varying gene tree estimation error (controlled by sequence length; 500-1500 bp) are shown in Figs. 3 (b). All methods showed improved accuracy with increasing sequence lengths, and best results were obtained on true gene trees. These results clearly show that wQFM-imputed is better than wQFM-incomplete and ASTRAL-complete and, in many cases, it is even better than wQFM-complete and ASTRAL-complete. Similar to 15-taxon dataset, we investigated the divergence of various quartet distributions from true quartet distributions (see Fig. 3(c)-(d)) and the quartet scores of different estimated species trees with respect to both estimated and true gene trees (see Section 4.2 in Supplementary Material) and observed similar trends, supporting our claim that QT-GILD effectively imputes missing data and accounts for gene tree estimation error while imputing the missing quartets.

### 4.3 Results on biological dataset

#### Angiosperm dataset

We re-analyzed the angriosperm (the most diverse plant clade) dataset from [53] containing 310 genes sampled from 42 angiosperms and 4 outgroups (three gymnosperms and one lycophyte). The goal is to resolve the position of *Amborella trichopoda Baill*. This dataset has a high level of missing data containing gene trees with as low as 16 taxa. On average, there are 12.9 missing taxa in the gene trees, resulting in a large number (30,129,163) of missing quartets. These missing quartets were imputed by QT-GILD. We estimated trees using ASTRAL and wQFM (with and without imputation). These three trees (Fig. 4) are highly congruent and placed *Amborella* as sister to water lilies (i.e., Nymphaeales) and rest of the angiosperms with high support. This relationship is consistent to the tree estimated by concatenation using maximum likelihood (reported in [53] as well as other molecular studies [46, 58–60]). However, alternative relationship (e.g., the placement of *Amborella* plus water lilies as sister to all other angiosperms) have also been reported [53,61,62]. wQFM-incomplete and wQFM-imputed recovered highly similar trees, differing only on a few branches with low support (e.g., the relative position of Fabales (*Glycine, Medicago*) and Fagales (*Betula, Quercus*) and the relationships within the clade containing *Musa, Phoenix, Oryza*, and *Sorghum*). The fact that there were a large proportion (71.33%) of missing quartets and QT-GILD imputes them by leveraging remaining 28.66% quartets, and yet the imputed distribution resulted in a meaningful angiosperm tree shows the efficacy of QT-GILD.

**Figure 4:**
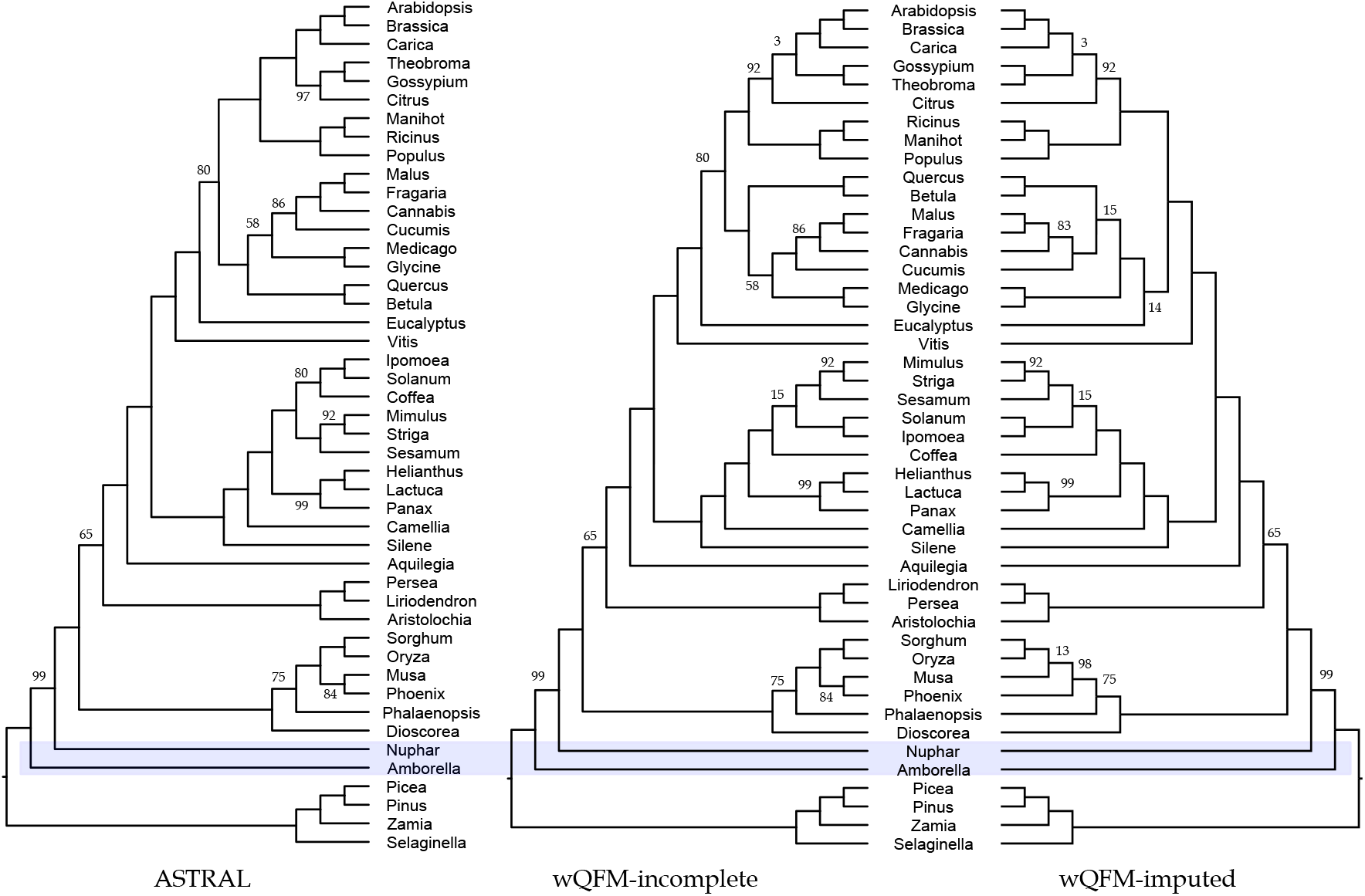
Analyses of the angiosperm dataset using ASTRAL and wQFM (with and without imputation). Branch supports (BS) represent quartet based local posterior probability [63] (multiplied by 100). All BS values are 100% except where noted.

### 4.4 Running time

The running time of QT-GILD mostly depends on the number of taxa, number of genes and the predefined number of epochs. We ran QT-GILD for 2000 epochs in our study. All analyses were run on the same machine with Intel Core i7-10700K CPU (16 cores), 64GB RAM, NVIDIA GeForce RTX 3070 GPU (8GB memory). For a single epoch, it takes 25 × 10^-3^ − 800 × 10^-3^ s on the 15-taxon dataset with varying numbers of genes. For 37-taxon datasets, it takes 1 − 6 s and, on the 46-taxon angiosperm dataset with 310 genes, QT-GILD takes 10−11 s to run (see Table S9 in Supplementary Material). The running time of QT-GILD is more sensitive to the number of taxa, as the size of quartet distributions grows exponentially with the number of taxa.

## 5 Conclusions

This study introduces the quartet distribution imputation problem and shows the power and feasibility of applying deep learning techniques for imputing quartet distributions. Our proposed QT-GILD not only imputes the missing quartets but it can also “correct” the estimated quartets – resulting in a set of quartets that is often closer to the quartet distribution of true gene trees than that of the estimated gene trees, and thereby accounting for gene tree estimation error. Experimental studies using both simulated and real biological dataset suggest that QT-GILD may result in dramatic improvements in species tree accuracy. Therefore, the idea of estimating species trees by imputing quartet distributions has merit and should be pursued and used in future phylogenomic studies. As an immediate extension of the current study we plan to evaluate QT-GILD on a diverse set of real biological datasets as the pattern of missing data is sufficiently complex and heterogeneous across various datasets. Automatic selection of various hyper parameters of this architecture is another important research avenue. Because QT-GILD tries to learn the overall quartet distribution guided by a self-supervised feedback loop and correct for gene tree estimation error, investigating its application beyond imputing incomplete gene trees where the goal would be to improve estimated gene tree distributions is another interesting direction to take. In this regard, we can mask a reasonably large number of quartets in the estimated complete quartet distribution and then impute them with QT-GILD, utilizing its error correcting utility. QT-GILD is currently not scalable to large datasets (both in terms of the number of taxa and genes), therefore this study was limited to small to moderate size datasets. Future study needs to develop a scalable variant of QT-GILD and assess its performance on large datasets.

## Supplementary Material

### 1 Overview

These supplementary materials present additional details about the quartet vector representation used by QT-GILD (Sec. 2) and the datasets analyzed in this study (Sec. 3), additional results on simulated and real biological datasets (Sec. 4), results including wQMC (Sec. 5), running time information (Sec. 6) and the values used for different hyper-parameters of QT-GILD (Sec. 7).

### 2 Quartet Encoding

**Figure S1:**
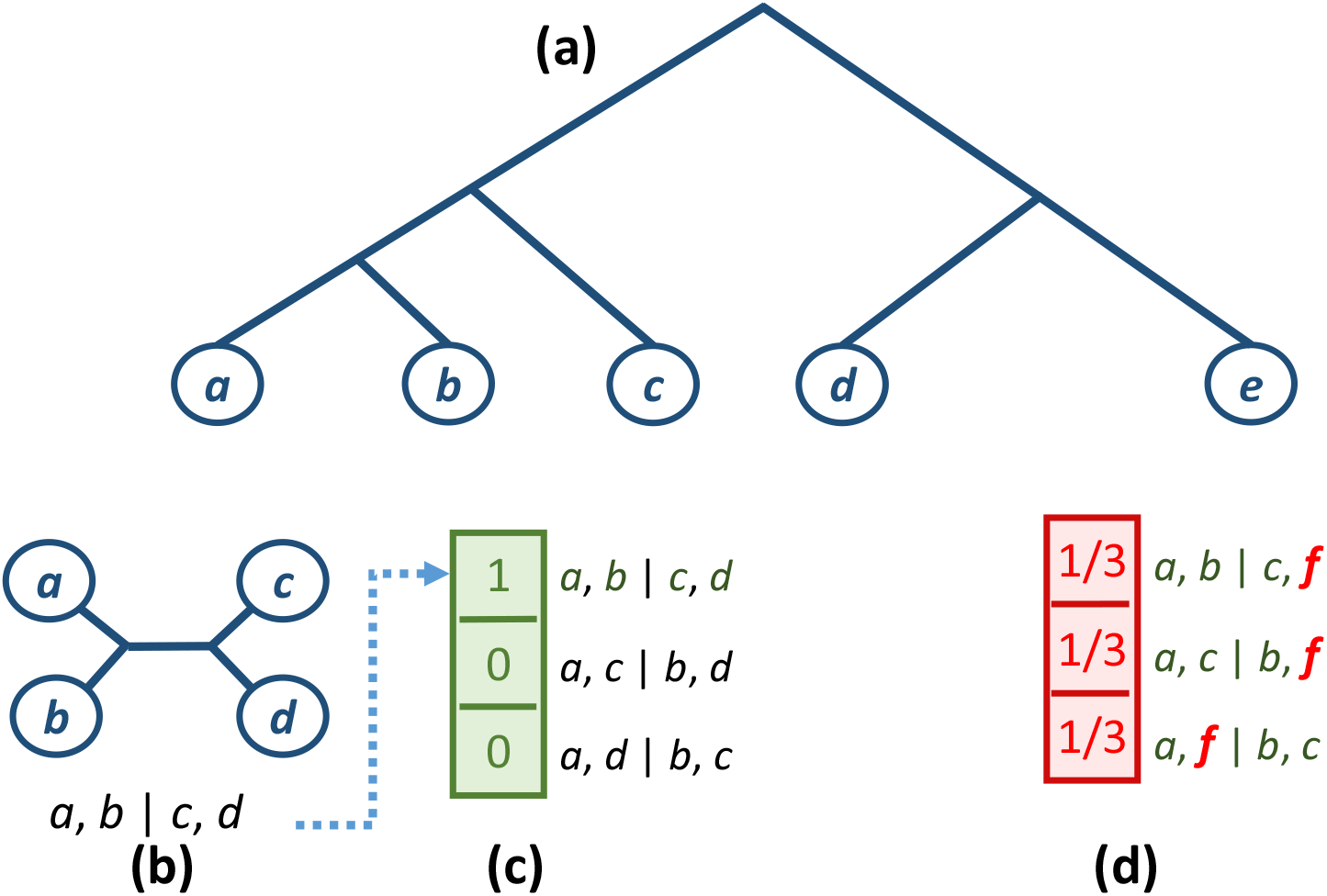
An example of *quartet vector* representation. (a) An incomplete gene tree (with respect to the full set of taxa *S* = {*a, b, c, d, e, f*}) in which a taxa *f* is missing. (b) The quartet *ab*|*cd*, on the set of four taxa {*a, b, c, d*}, which is present in the gene tree. (c) *Quartet vector* representation for the set of four taxa {*a,b,c,d*}. (d) *Quartet vector* representation for the four taxa set {*a,b,c,f*}, where the probabilities of all possible quartets are the same (i.e., 1/3) since these quartets are missing in the gene tree.

### 3 Dataset

**Table S1:**
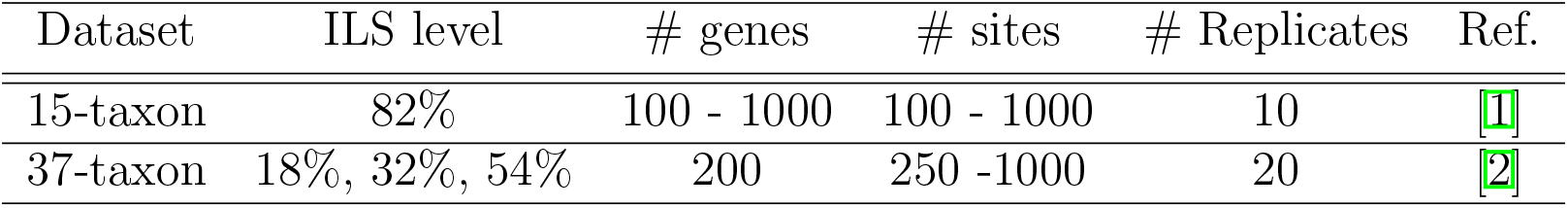
Properties of the simulated datasets. Level of ILS is presented in terms of the average topological distance between true gene trees and true species tree.

### 4 Additional Results

#### 4.1 Results on 15-taxon dataset

**Table S2:** Quartet statistics for 15-taxon dataset. For various model conditions with different numbers of missing taxa, we show the average number of quartets (present and missing) over 10 replicates. We also show what proportion of the missing quartets that were correctly imputed by QT-GILD with respect to both estimated and true gene trees.

| Model | # of Missing Taxa | # of Quartets | # of Missing Quartets | % Missing Quartets | % Correctly imputed (w.r.t. estimated gene trees) | % Correctly imputed (w.r.t. true gene trees) |
| --- | --- | --- | --- | --- | --- | --- |
| 100gene-100bp | 0 | 136500 | 0.0 | 0.0 | — | — |
|  | 1-2 | 118227.2 | 18272.8 | 13.39 | 54.67 | 52.20 |
|  | 2-3 | 85288.8 | 51211.2 | 37.5 | 54.40 | 52.68 |
|  | 4-5 | 41301.2 | 95198.8 | 69.74 | 51.73 | 52.23 |
|  | 5-6 | 27046 | 109454 | 80 | 51.81 | 51.03 |
| 100gene-1000bp | 0 | 136500 | 0.0 | 0.0 | — | — |
|  | 1-2 | 118119 | 18381 | 13.46 | 55.72 | 63.18 |
|  | 2-3 | 85292 | 51207 | 37.5 | 54.96 | 62.72 |
|  | 4-5 | 41291 | 95209 | 69.7 | 54.30 | 60.03 |
|  | 5-6 | 27503 | 108897 | 79.85 | 53.44 | 58.56 |
| 1000gene-100bp | 0 | 1365000 | 0.0 | 0.0 | — | — |
|  | 1-2 | 1183509.6 | 181490.4 | 13.29 | 54.92 | 56.28 |
|  | 2-3 | 858228.8 | 506771.2 | 37.12 | 54.42 | 56.34 |
|  | 4-5 | 411790.5 | 953209.5 | 69.8 | 53.53 | 54.26 |
|  | 5-6 | 269976 | 1095024 | 80.2 | 52.47 | 53.21 |
| 1000gene-1000bp | 0 | 1365000 | 0.0 | 0.0 | — | — |
|  | 1-2 | 1181069.8 | 183930.2 | 13.47 | 58.04 | 62.88 |
|  | 2-3 | 857113.4 | 507886.6 | 37.2 | 57.81 | 62.76 |
|  | 4-5 | 411361.5 | 953638.5 | 69.8 | 56.44 | 62.02 |
|  | 5-6 | 270435 | 1094565 | 80.18 | 55.38 | 60.65 |

**Figure S2:**
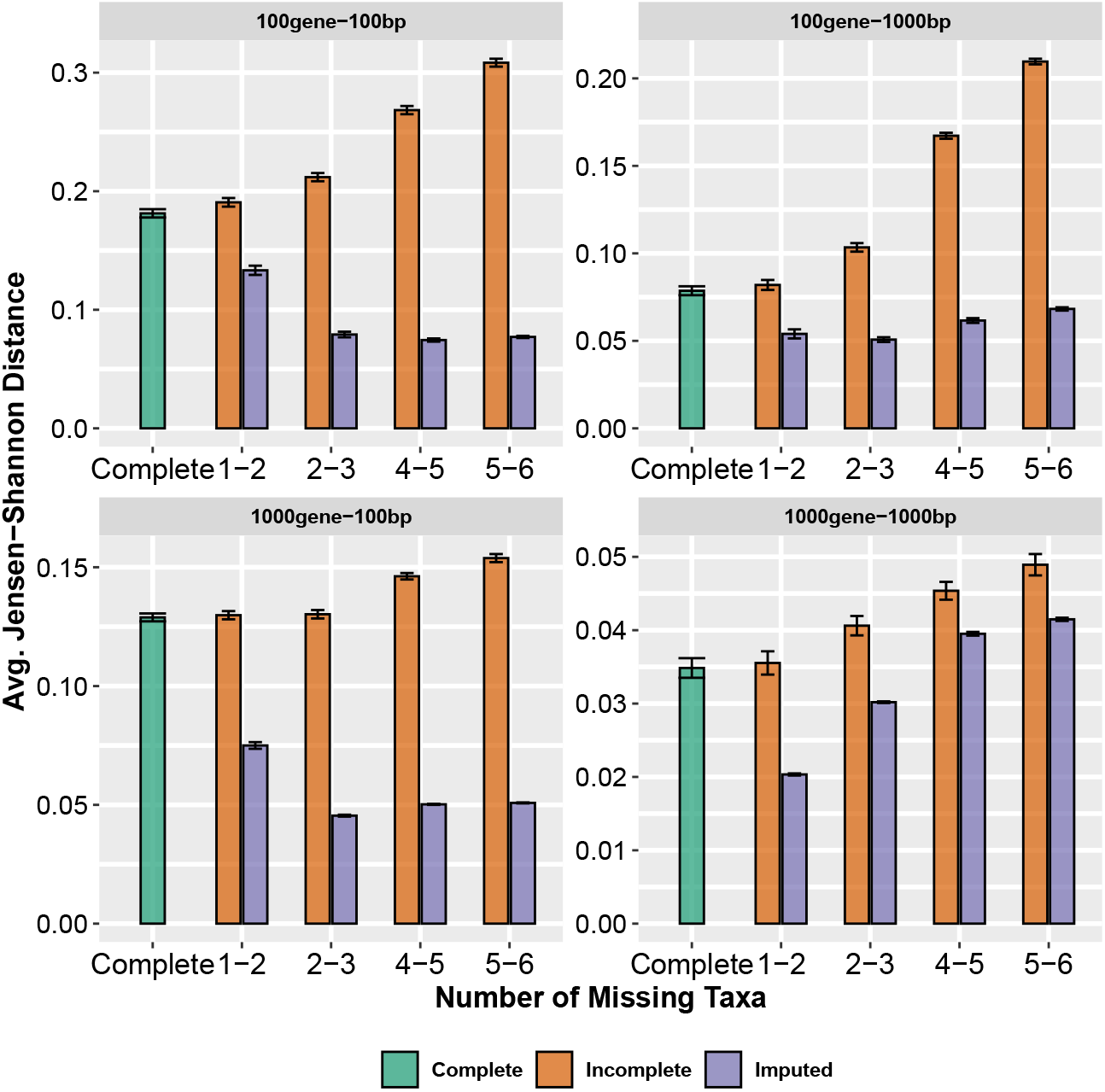
Jansen-Shannon divergence between true quartet distributions and complete, incomplete and imputed quartet distributions. For each model condition, we varied the taxa removal rate from ~6% to ~40%.

**Table S3:** Average quartet scores of the estimated and true species tree with respect to estimated gene trees for the 15-Taxon dataset. The quartet scores that are equal to the true species tree score are shown in bold and the highest quartet scores are shown in italic.

| Model | Number of Missing Taxa | Complete (Astral) | Complete (WQFM) | Incomplete (Astral) | Incomplete (WQFM) | Imputed (WQFM) | True Species Tree |
| --- | --- | --- | --- | --- | --- | --- | --- |
| 100gene-100bp | 0 | *69933.8* | 69776.6 | — | — | — | 69307.4 |
|  | 1-2 | — | — | *69914.8* | 69784.1 | 69678.7 |  |
|  | 2-3 | — | — | *69885.2* | 69632.9 | 69320.8 |  |
|  | 4-5 | — | — | *69559.1* | 69366.6 | **69307.4** |  |
|  | 5-6 | — | — | 69307.8 | 69179.2 | ***69307.4*** |  |
| 100gene-1000bp | 0 | *82166.6* | 82109.2 | — | — | — | 82099.2 |
|  | 1-2 | — | — | *82166.6* | 82146.4 | **82099.2** |  |
|  | 2-3 | — | — | *82106* | 82083 | **82099.2** |  |
|  | 4-5 | — | — | 81998.1 | 82022.5 | ***82099.2*** |  |
|  | 5-6 | — | — | 81682.7 | 81761.3 | ***82099.2*** |  |
| 1000gene-100bp | 0 | *693656.1* | 692949.2 | — | — | — | 690268 |
|  | 1-2 | — | — | *693648.1* | 693172.4 | **690268** |  |
|  | 2-3 | — | — | *693615.8* | 693008.6 | **690268** |  |
|  | 4-5 | — | — | *693533.8* | 693117.7 | **690268** |  |
|  | 5-6 | — | — | *693527.5* | 692743.1 | **690268** |  |
| 1000gene-1000bp | 0 | *818022* | 817994 | — | — | — | 817937.3 |
|  | 1-2 | — | — | *818021* | 8187994 | **817937.3** |  |
|  | 2-3 | — | — | *818003.9* | 817993.9 | **817937.3** |  |
|  | 4-5 | — | — | *818022.2* | 817993.9 | **817937.3** |  |
|  | 5-6 | — | — | *817994* | 817993.9 | **817937.3** |  |

**Table S4:** Average quartet scores (over 10 replicates) of the estimated and true species trees with respect to true gene trees for the 15-Taxon dataset. The quartet scores that are equal to the true species tree score are shown in bold and the highest quartet scores are shown in italic.

| Model | Number of Missing Taxa | Complete (Astral) | Complete (WQFM) | Incomplete (Astral) | Incomplete (WQFM) | Imputed (WQFM) | True species Tree |
| --- | --- | --- | --- | --- | --- | --- | --- |
| 100gene-100bp | 0 | 82158.6 | *82708.3* | — | — | — | 84634.9 |
|  | 1-2 | — | — | 81790.7 | 82713.1 | *83549.3* |  |
|  | 2-3 | — | — | 81947.2 | 82231.6 | *84627.5* |  |
|  | 4-5 | — | — | 81213.6 | 80831.3 | ***84634.9*** |  |
|  | 5-6 | — | — | 80699.4 | 80624 | ***84634.9*** |  |
| 100gene-1000bp | 0 | 84345.1 | *84366.1* | — | — | — | 84634.9 |
|  | 1-2 | — | — | 82166.6 | 82146.4 | ***84634.9*** |  |
|  | 2-3 | — | — | 82106 | 82083 | ***84634.9*** |  |
|  | 4-5 | — | — | 81998.1 | 82022.5 | ***84634.9*** |  |
|  | 5-6 | — | — | 81682.7 | 81761.3 | ***84634.9*** |  |
| 1000gene-100bp | 0 | 827970.1 | *834890.2* | — | — | — | 844184.3 |
|  | 1-2 | — | — | 826981.3 | 833742.8 | ***844184.3*** |  |
|  | 2-3 | — | — | 825679.5 | 832472.8 | ***844184.3*** |  |
|  | 4-5 | — | — | 824174.4 | 830065.4 | ***844184.3*** |  |
|  | 5-6 | — | — | 824202.9 | 831620.6 | ***844184.3*** |  |
| 1000gene-1000bp | 0 | 843362.8 | *843791.8* | — | — | — | 844184.3 |
|  | 1-2 | — | — | 818021 | 8187994 | ***844184.3*** |  |
|  | 2-3 | — | — | 842977 | 843791.8 | ***844184.3*** |  |
|  | 4-5 | — | — | 843362.8 | 843791.8 | ***844184.3*** |  |
|  | 5-6 | — | — | 843162.5 | 843791.8 | ***844184.3*** |  |

#### 4.2 Additional results on 37-taxon dataset

**Table S5:** Quartet statistics for 37-taxon dataset. For varying levels of ILS and different numbers of missing taxa, we show the number of quartets (present and missing). We also show what proportion of the missing taxa were correctly imputed by QT-GILD.

| Model | # of Missing Taxa | # of Quartets | # of Missing Quartets | % Missing Quartets |
| --- | --- | --- | --- | --- |
| 0.5X | 0 | 13209000 | 0.0 | — |
|  | 1-2 | 12476793 | 732207 | 6 |
|  | 2-3 | 11131667 | 2077333 | 16 |
|  | 3-4 | 9886466 | 3322534 | 25 |
| 1X | 0 | 13209000 | 0.0 | — |
|  | 1-2 | 12492223 | 716777 | 5.42 |
|  | 2-3 | 11144980 | 2064020 | 16 |
|  | 3-4 | 9877714 | 3331286 | 25 |
| 2X | 0 | 13209000 | 0.0 | — |
|  | 1-2 | 12491994 | 717006 | 5 |
|  | 2-3 | 11113410 | 2095590 | 16 |
|  | 3-4 | 9877257 | 3331743 | 25 |

**Table S6:**
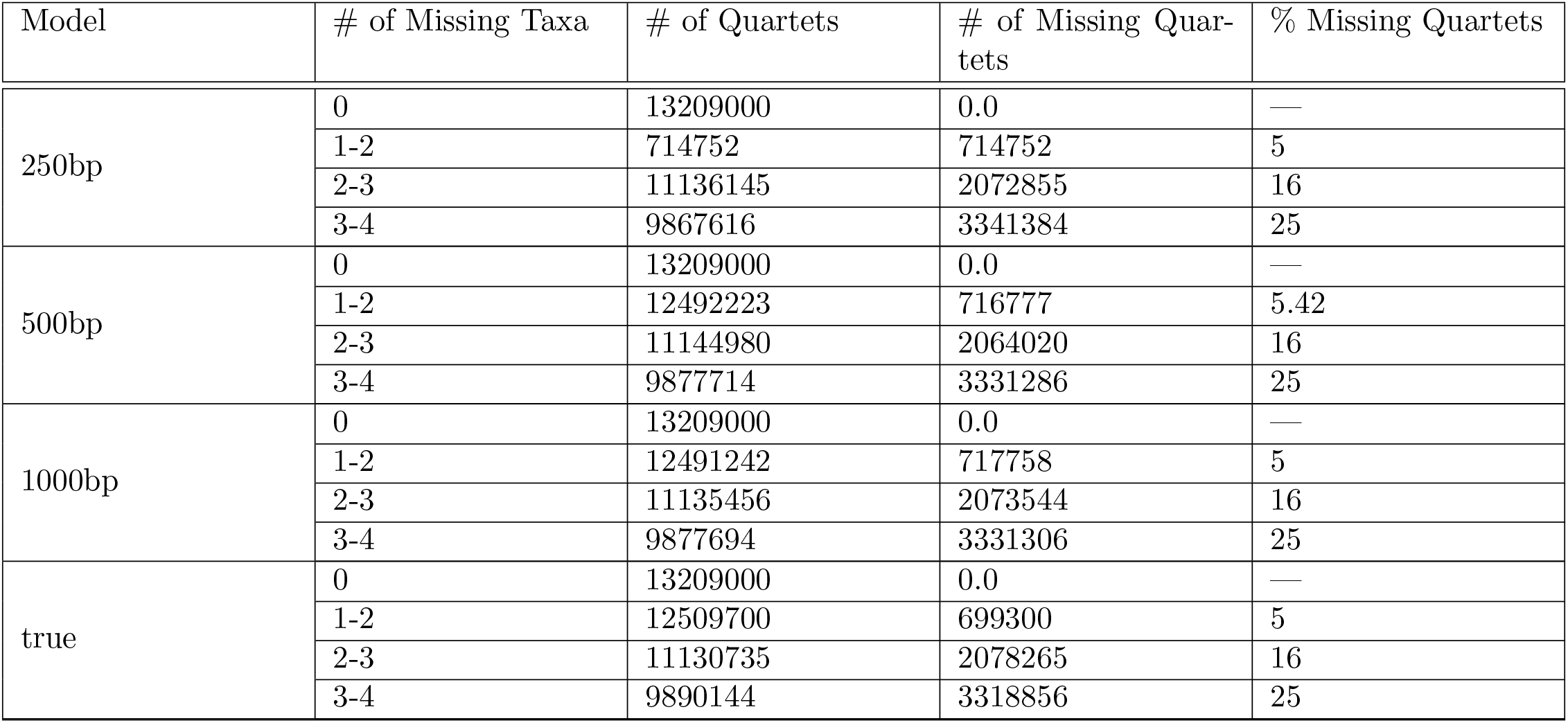
Quartet statistics for 37-taxon dataset. For varying amounts of gene tree estimation errors (controlled by sequence length) and different numbers of missing taxa, we show the number of quartets (present and missing). We also show what proportion of the missing taxa were correctly imputed by QT-GILD.

| Model | # of Missing Taxa | # of Quartets | # of Missing Quartets | % Missing Quartets |
| --- | --- | --- | --- | --- |
| 250bp | 0 | 13209000 | 0.0 | — |
|  | 1-2 | 714752 | 714752 | 5 |
|  | 2-3 | 11136145 | 2072855 | 16 |
|  | 3-4 | 9867616 | 3341384 | 25 |
| 500bp | 0 | 13209000 | 0.0 | — |
|  | 1-2 | 12492223 | 716777 | 5.42 |
|  | 2-3 | 11144980 | 2064020 | 16 |
|  | 3-4 | 9877714 | 3331286 | 25 |
| 1000bp | 0 | 13209000 | 0.0 | — |
|  | 1-2 | 12491242 | 717758 | 5 |
|  | 2-3 | 11135456 | 2073544 | 16 |
|  | 3-4 | 9877694 | 3331306 | 25 |
| true | 0 | 13209000 | 0.0 | — |
|  | 1-2 | 12509700 | 699300 | 5 |
|  | 2-3 | 11130735 | 2078265 | 16 |
|  | 3-4 | 9890144 | 3318856 | 25 |

**Table S7:** Average quartet scores (over 20 replicates) of the estimated and true species trees with respect to estimated gene trees for 37-Taxon datasets. The highest quartet scores are shown in italic.

| Model | Number of Missing Taxa | Complete (Astral) | Complete (WQFM) | Incomplete (Astral) | Incomplete (WQFM) | Imputed (WQFM) | True |
| --- | --- | --- | --- | --- | --- | --- | --- |
| 0.5X | 0 | *10006308.5* | 10005483.8 | — | — | — | 10002171.5 |
|  | 1-2 | — | — | *10006308.5* | 10005684.3 | 10005787.4 |  |
|  | 2-3 | — | — | *10005758.1* | 10004869.1 | 10004204 |  |
|  | 3-4 | — | — | *10004950* | 10004453.5 | 10003461 |  |
| 1X | 0 | *11262866* | 11262373 | — | — | — | 11260000 |
|  | 1-2 | — | — | *11262866* | 11262649.5 | 11259262 |  |
|  | 2-3 | — | — | *11262758.5* | 11262373.75 | 11262247.9 |  |
|  | 3-4 | — | — | *1126207* | 11261781 | 11258958 |  |
| 2X | 0 | *11950022* | 11949736 | — | — | — | 11949183 |
|  | 1-2 | — | — | *11950014.5* | 11949729 | 11949599.5 |  |
|  | 2-3 | — | — | *11949893.2* | 11949750.2 | 11949525.1 |  |
|  | 3-4 | — | — | *11949882.2* | 11949510 | 11948202.6 |  |

**Table S8:** Average quartet scores (over 20 replicates) of the estimated species trees and the true species tree with respect to true gene trees for 37-Taxon datasets. The highest quartet scores are shown in italic.

| Model | Number of Missing Taxa | Complete (Astral) | Complete (WQFM) | Incomplete (Astral) | Incomplete (WQFM) | Imputed (WQFM) | True |
| --- | --- | --- | --- | --- | --- | --- | --- |
| 0.5X | 0 | 11717528.6 | *11719898.1* | — | — | — | 11735843.2 |
|  | 1-2 | — | — | 11717528.6 | 11720236.7 | *11721154.4* |  |
|  | 2-3 | — | — | 11713398.2 | 11712811.8 | *11713814.4* |  |
|  | 3-4 | — | — | 11709382.1 | 11709370.5 | *11710978.9* |  |
| 1X | 0 | 11730467.8 | *11730710* | — | — | — | 11736367.4 |
|  | 1-2 | — | — | 11731675.3 | 11731322.85 | *11734827.2* |  |
|  | 2-3 | — | — | 11729895 | 11730710.6 | *11731448.8* |  |
|  | 3-4 | — | — | 11727661.7 | 11730502.4 | *11733830* |  |
| 2X | 0 | 11745806.42 | *11747526.3* | — | — | — | 11748585.4 |
|  | 1-2 | — | — | 11745484 | *11747408.2* | 11746526.3 |  |
|  | 2-3 | — | — | 11746648 | *11747028.7* | 11745633.6 |  |
|  | 3-4 | — | — | 11745714 | *11747301.85* | 11742943.9 |  |

### 5 Results with wQMC

We present results that include wQMC, so the figures shown here include ASTRAL, wQFM, and wQMC. These results, which include two quartet amalgamation techniques (wQMC and wQFM), help to assess the impact of imputed quartet distributions (produced by QT-GILD) on different quartet amalgamation techniques.

#### 5.1 Results on 15-taxon dataset

**Figure S3:**
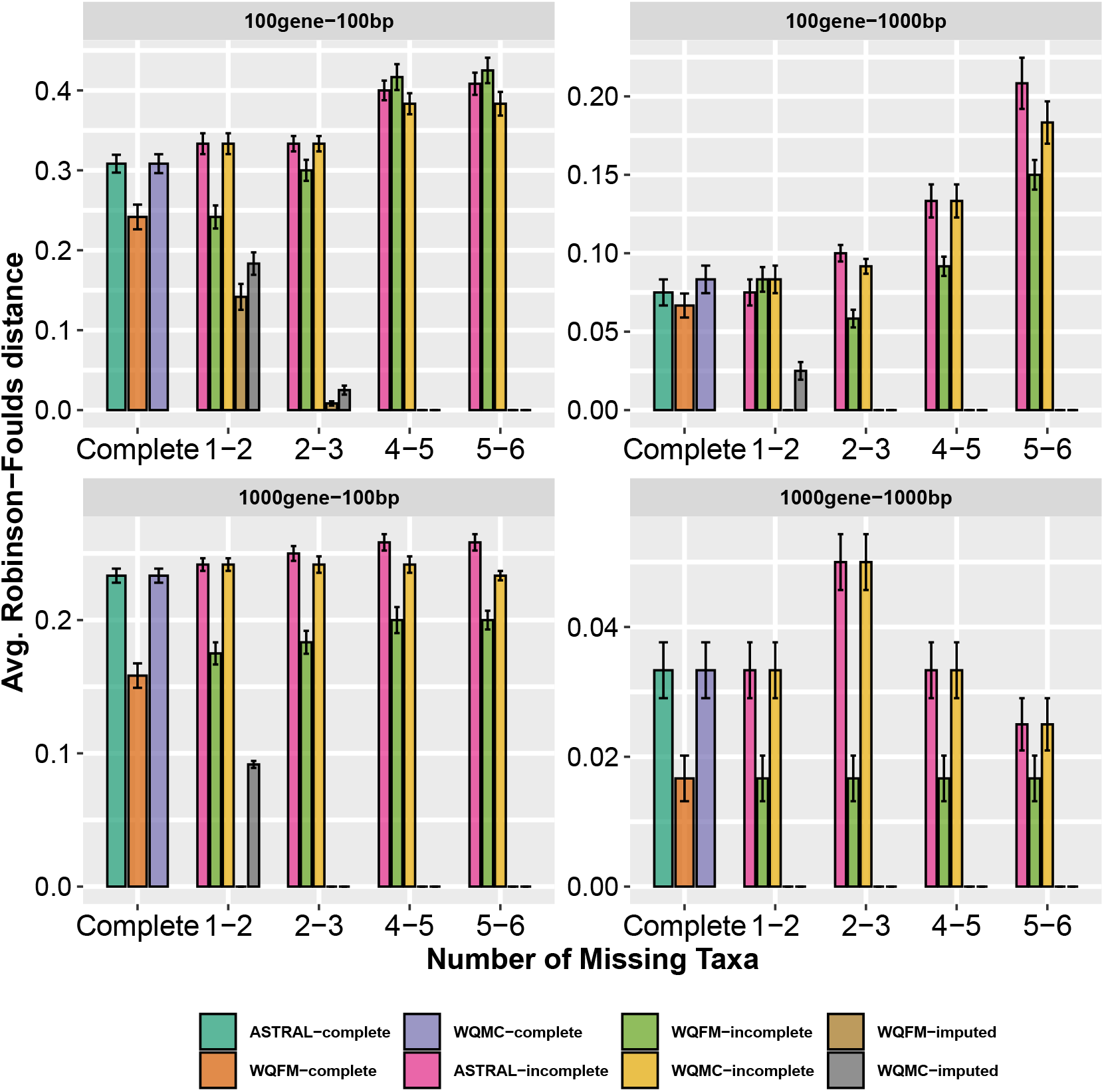
Comparison of different variants of ASTRAL, wQMC and wQFM on 15-taxon dataset. We show the average RF rates with standard errors over 10 replicates. For each model condition, we varied the taxa removal rate from ~6% to ~40%, resulting in 13-80% missing quartets.

#### 5.2 Results on 37-taxon dataset

**Figure S4:**
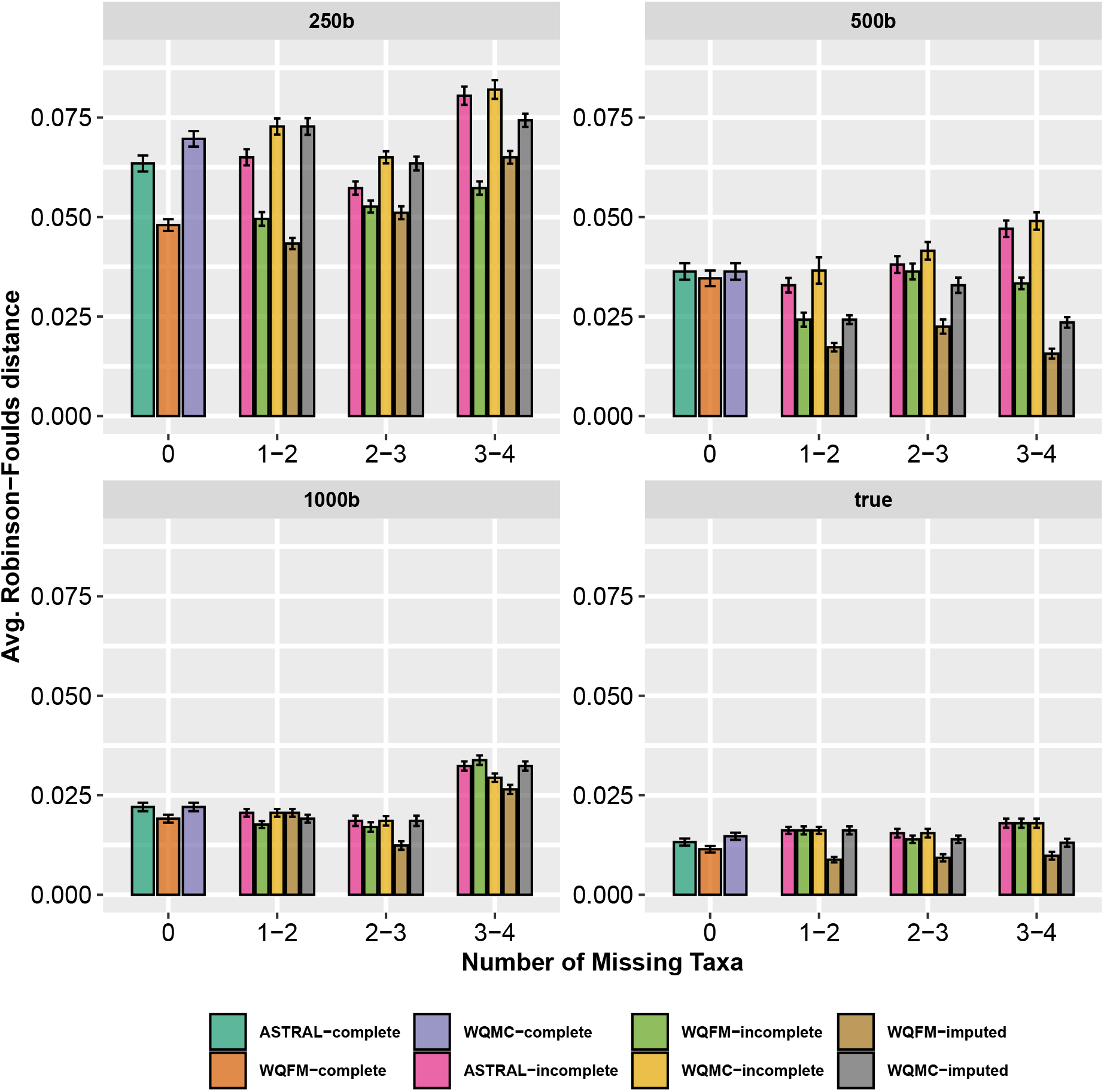
Comparison of different variants of ASTRAL, wQMC and wQFM on 37-taxon dataset with varying amounts of gene tree estimation error. We show the average RF rates of 37-taxon dataset with standard errors over 20 replicates. The sequence length was varied from 250bp to 1500bp, keeping the number of genes fixed at 200, and ILS at 1X (moderate ILS). We also analyzed the true gene trees.

**Figure S5:**
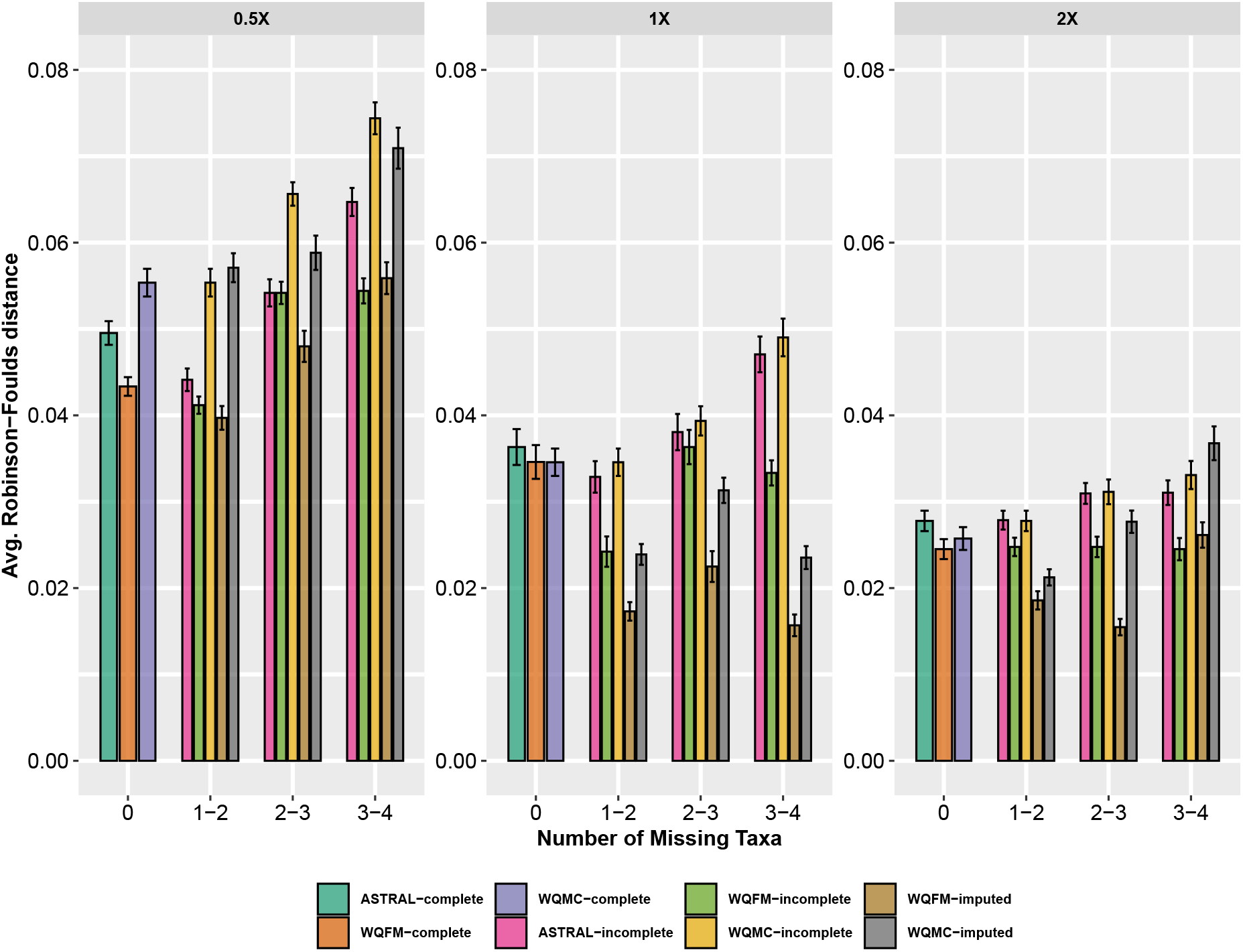
Comparison of different variants of ASTRAL, wQMC and wQFM on 37-taxon dataset with varying levels of gene tree discordance. We show the average RF rates of 37-taxon dataset with standard errors over 20 replicates. The level of ILS was varied from 0.5X (highest) to 2X (lowest) amount, keeping the sequence length fixed at 500bp and the number of genes at 200.

#### 5.3 Results on biological dataset (Angiosperm dataset)

wQMC estimated trees are congruent to the trees estimated by ASTRAL and wQFM and placed *Amborella* as sister to water lilies (i.e., Nymphaeales) and rest of the angiosperms with high support. wQMC-incomplete and wQMC-imputed recovered highly similar trees, differing only on a few branches with low support.

**Figure S6:**
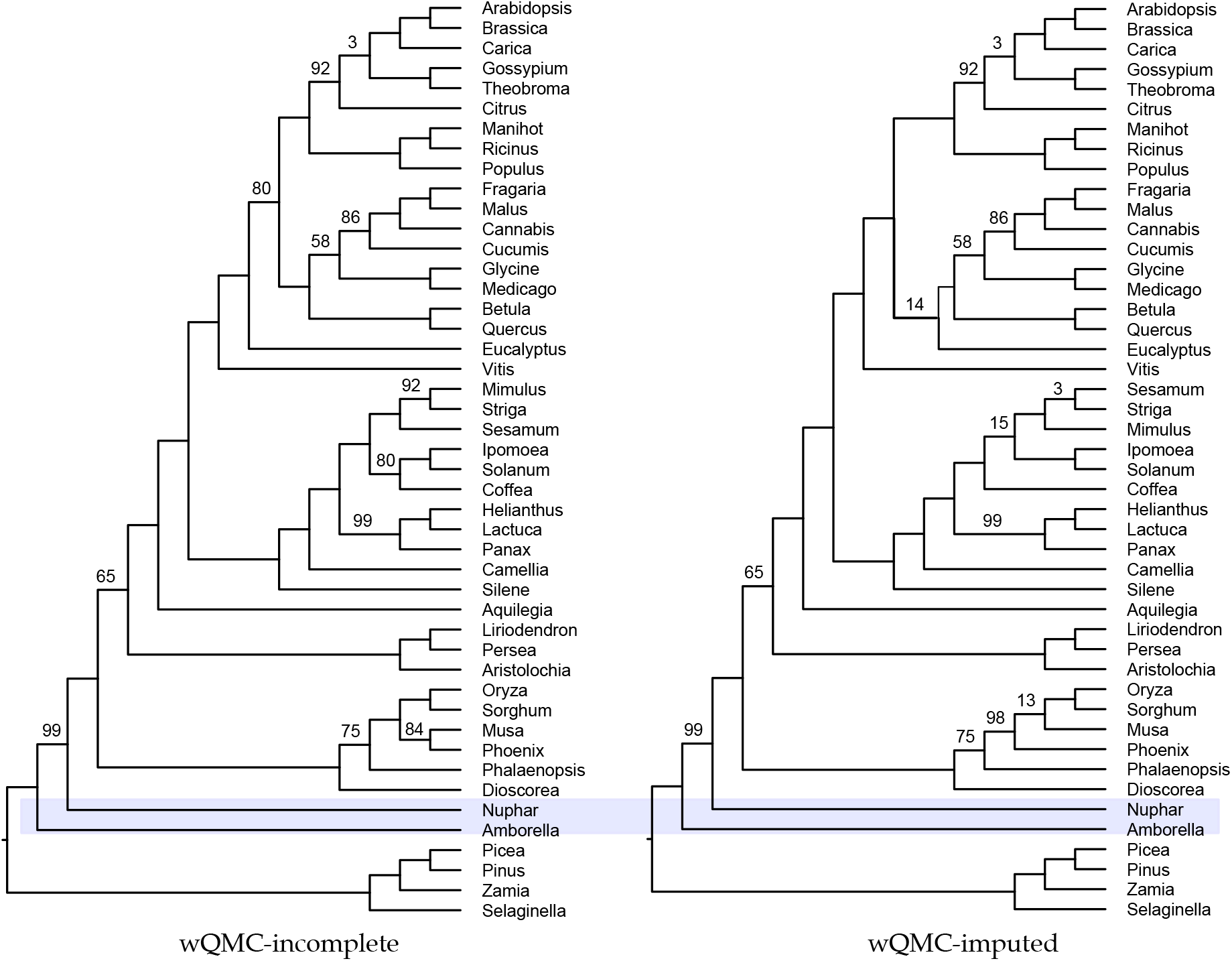
Analyses of the angiosperm dataset using wQMC (with and without imputation). Branch supports (BS) represent quartet based local posterior probability [3] (multiplied by 100). All BS values are 100% except where noted.

### 6 Running time

**Table S9:** Running time of QT-GILD for one epoch on different dataset. Across all the datasets, we ran QT-GILD for 2000 epochs.

| Dataset | # Genes | Running time (seconds) |
| --- | --- | --- |
| 15-taxon | 100 | $25 \times 10^{-3} - 26 \times 10^{-3}$ |
| 15-taxon | 1000 | $750 \times 10^{-3} - 800 \times 10^{-3}$ |
| 37-taxon | 100 | 1 – 1.5 |
| 37-taxon | 200 | 2 – 3 |
| 37-taxon | 400 | 5 – 6 |
| Angiosperm (46-taxon) | 310 | 10 – 11 |

### 7 Hyper-parameters of QT-GILD

**Table S10:**
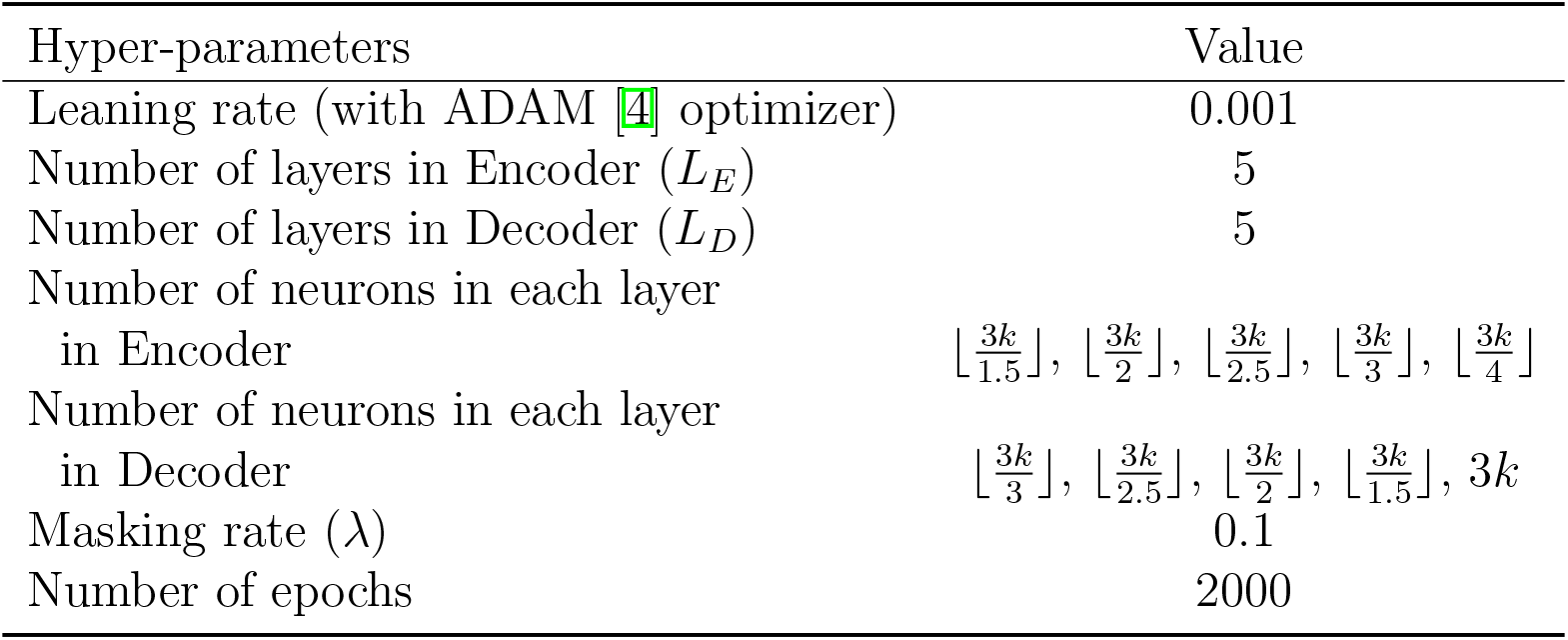
Various hyper-parameters and their values used in our experiments. Here *k* is the number of gene-trees.

| Hyper-parameters | Value |
| --- | --- |
| Learning rate (with ADAM [4] optimizer) | 0.001 |
| Number of layers in Encoder ($L_E$) | 5 |
| Number of layers in Decoder ($L_D$) | 5 |
| Number of neurons in each layer<br>in Encoder | $\lfloor \frac{3k}{1.5} \rfloor, \lfloor \frac{3k}{2} \rfloor, \lfloor \frac{3k}{2.5} \rfloor, \lfloor \frac{3k}{3} \rfloor, \lfloor \frac{3k}{4} \rfloor$ |
| Number of neurons in each layer<br>in Decoder | $\lfloor \frac{3k}{3} \rfloor, \lfloor \frac{3k}{2.5} \rfloor, \lfloor \frac{3k}{2} \rfloor, \lfloor \frac{3k}{1.5} \rfloor, 3k$ |
| Masking rate ($\lambda$) | 0.1 |
| Number of epochs | 2000 |

## References

[1] W. P. Maddison. Gene trees in species trees. Systematic Biology, 46:523–536, 1997.

[2] J H Degnan and N A Rosenberg. Discordance of species trees with their most likely gene trees. PLoS Genetics, 2:762 – 768, 2006.

[3] J. H. Degnan and L. A. Salter. Gene tree distributions under the coalescent process. Evolution, 59(1):24–37, January 2005.

[4] R. R. Hudson. Testing the constant-rate neutral allele model with protein sequence data. Evolution, 37:203 – 217, 1983.

[5] M. Nei. Stochastic errors in DNA evolution and molecular phylogeny. In H. Gershowitz, D. L. Rucknagel, and R. E. Tashian, editors, Evolutionary Perspectives and the New Genetics, pages 133 – 147, 1986.

[6] M. Nei. Molecular evolutionary genetics. New York, 1987. Columbia University Press.

[7] N. Rosenberg. The Probability of Topological Concordance of Gene Trees and Species Trees. Theoretical Population Biology, 61(2):225–247, March 2002.

[8] Fumio Tajima. Evolutionary relationship of DNA sequences in finite populations. Genetics, 105(2):437–460, October 1983.

[9] N. Takahata. Gene geneaology in three related populations: consistency probability between gene and population trees. Genetics, 122:957–966, 1989.

[10] Sebastien Roch and Mike Steel. Likelihood-based tree reconstruction on a concatenation of aligned sequence data sets can be statistically inconsistent. Theoretical Population Biology, 100:56–62, 2015.

[11] J H Degnan, M DeGiorgio, D Bryant, and N A Rosenberg. Properties of consensus methods for inferring species trees from gene trees. Systematic Biology, 58:35–54, 2009.

[12] L S Kubatko and J H Degnan. Inconsistency of phylogenetic estimates from concatenated data under coalescence. Systematic Biology, 56:17, 2007.

[13] Liang Liu, Lili Yu, and Scott V Edwards. A maximum pseudo-likelihood approach for estimating species trees under the coalescent model. BMC Evolutionary Biology, 10:302, 2010.

[14] Siavash Mirarab, Rezwana Reaz, Md S Bayzid, Théo Zimmermann, M Shel Swenson, and Tandy Warnow. ASTRAL: genome-scale coalescent-based species tree estimation. Bioinformatics, 30(17):i541–i548, 2014.

[15] B Larget, S K Kotha, C N Dewey, and C Ané. BUCKy: Gene tree/species tree reconciliation with the Bayesian concordance analysis. Bioinformatics, 26(22):2910–2911, 2010.

[16] E Mossel and S Roch. Incomplete lineage sorting: consistent phylogeny estimation from multiple loci. IEEE/ACM Transactions on Computational Biology and Bioinformatics, 7(1):166–171, 2011.

[17] L S Kubatko, B C Carstens, and L L Knowles. Stem: Species tree estimation using maximum likelihood for gene trees under coalescence. Bioinformatics, 25:971–973, 2009.

[18] Julia Chifman and Laura Kubatko. Quartet from SNP data under the coalescent model. Bioinformatics, 30(23):3317–3324, 2014.

[19] Liang Liu, Lili Yu, Dennis K Pearl, and Scott V Edwards. Estimating species phylogenies using coalescence times among sequences. Systematic Biology, 58(5):468–477, 2009.

[20] Liang Liu and Lili Yu. Estimating species trees from unrooted gene trees. Systematic Biology, 60(5):661–667, 2011.

[21] Pranjal Vachaspati and Tandy Warnow. Astrid: accurate species trees from internode distances. BMC Genomics, 16(10):S3, 2015.

[22] Mazharul Islam, Kowshika Sarker, Trisha Das, Rezwana Reaz, and Md Shamsuzzoha Bayzid. Stelar: A statistically consistent coalescent-based species tree estimation method by maximizing triplet consistency. BMC Genomics, 21(1):1–13, 2020.

[23] L Liu. BEST: Bayesian estimation of species trees under the coalescent model. Bioinformatics, 24:2542–2543, 2008.

[24] J Heled and A J Drummond. Bayesian inference of species trees from multilocus data. Molecular Biology and Evolution, 27:570–580, 2010.

[25] Md Shamsuzzoha Bayzid and Tandy Warnow. Naive binning improves phylogenomic analyses. Bioinformatics, 29(18):2277–2284, 2013.

[26] Brian Tilston Smith, Michael G Harvey, Brant C Faircloth, Travis C Glenn, and Robb T Brumfield. Target capture and massively parallel sequencing of ultraconserved elements for comparative studies at shallow evolutionary time scales. Systematic Biology, 63(1):83–95, 2013.

[27] Md Shamsuzzoha Bayzid and Tandy Warnow. Estimating optimal species trees from incomplete gene trees under deep coalescence. Journal of Computational Biology, 19(6):591–605, 2012.

[28] A D Leaché and B Rannala. The accuracy of species tree estimation under simulation: a comparison of methods. Systematic Biology, 60(2):126–137, 2011.

[29] Peter A Hosner, Brant C Faircloth, Travis C Glenn, Edward L Braun, and Rebecca T Kimball. Avoiding missing data biases in phylogenomic inference: an empirical study in the landfowl (aves: Galliformes). Molecular Biology and Evolution, 33(4):1110–1125, 2016.

[30] Jeffrey W Streicher, James A Schulte, and John J Wiens. How should genes and taxa be sampled for phylogenomic analyses with missing data? an empirical study in iguanian lizards. Systematic Biology, 65(1):128–145, 2016.

[31] Zhenxiang Xi, Liang Liu, and Charles C Davis. The impact of missing data on species tree estimation. Molecular Biology and Evolution, 33(3):838–860, 2016.

[32] M. S. Bayzid and T. Warnow. Gene tree parsimony for incomplete gene trees: addressing true biological loss. Algorithms for Molecular Biology, 13:1, 2018.

[33] Alan R Lemmon, Jeremy M Brown, Kathrin Stanger-Hall, and Emily Moriarty Lemmon. The effect of ambiguous data on phylogenetic estimates obtained by maximum likelihood and bayesian inference. Systematic biology, 58(1):130–145, 2009.

[34] J Gordon Burleigh, Khidir W Hilu, and Douglas E Soltis. Inferring phylogenies with incomplete data sets: a 5-gene, 567-taxon analysis of angiosperms. BMC evolutionary biology, 9(1):1–11, 2009.

[35] Michael Nute, Jed Chou, Erin K Molloy, and Tandy Warnow. The performance of coalescent-based species tree estimation methods under models of missing data. BMC Genomics, 19(5):1–22, 2018.

[36] Michael J Sanderson, Michelle M McMahon, and Mike Steel. Terraces in phylogenetic tree space. Science, 333(6041):448–450, 2011.

[37] Ishrat Tanzila Farah, Muktadirul Islam, Kazi Tasnim Zinat, Atif Hasan Rahman, and Shamsuzzoha Bayzid. Species tree estimation from gene trees by minimizing deep coalescence and maximizing quartet consistency: A comparative study and the presence of pseudo species tree terraces. Systematic Biology, 70(6):1213–1231, 04 2021.

[38] J H Degnan and N A Rosenberg. Gene tree discordance, phylogenetic inference and the multispecies coalescent. Trends in Ecology and Evolution, 26(6), 2009.

[39] James H Degnan. Anomalous unrooted gene trees. Systematic Biology, 62(4):574–590, 2013.

[40] K Strimmer and A von Haeseler. Quartet puzzling: A quartet maximim-likelihood method for reconstructing tree topologies. Molecular Biology and Evolution, 13(7):964–969, 1996.

[41] Heiko A Schmidt, Korbinian Strimmer, Martin Vingron, and Arndt von Haeseler. Tree-puzzle: maximum likelihood phylogenetic analysis using quartets and parallel computing. Bioinformatics, 18(3):502–504, 2002.

[42] Vincent Ranwez and Olivier Gascuel. Quartet-based phylogenetic inference: improvements and limits. Molecular Biology and Evolution, 18(6):1103–1116, 2001.

[43] S. Snir and S. Rao. Quartets MaxCut: a divide and conquer quartets algorithm. IEEE/ACM Transactions on Computational Biology and Bioinformatics, 7(4):704–718, 2010.

[44] Rezwana Reaz, Md Shamsuzzoha Bayzid, and M Sohel Rahman. Accurate phylogenetic tree reconstruction from quartets: A heuristic approach. PLoS ONE, 9(8):e104008, 2014.

[45] Eliran Avni, Reuven Cohen, and Sagi Snir. Weighted quartets phylogenetics. Systematic Biology, 64(2):233–242, 2015.

[46] Mahim Mahbub, Zahin Wahab, Rezwana Reaz, M Saifur Rahman, and Md Shamsuzzoha Bayzid. wQFM: highly accurate genome-scale species tree estimation from weighted quartets. Bioinformatics, 06 2021.

[47] Sarah Christensen, Erin K Molloy, Pranjal Vachaspati, and Tandy Warnow. Octal: Optimal completion of gene trees in polynomial time. Algorithms for Molecular Biology, 13(1):1–18, 2018.

[48] Ashish Vaswani, Noam Shazeer, Niki Parmar, Jakob Uszkoreit, Llion Jones, Aidan N Gomez, Lukasz Kaiser, and Illia Polosukhin. Attention is all you need. In NIPS, 2017.

[49] Mostofa Rafid Uddin, Sazan Mahbub, M Saifur Rahman, and Md Shamsuzzoha Bayzid. SAINT: self-attention augmented inception-inside-inception network improves protein secondary structure prediction. Bioinformatics, 36(17):4599–4608, 2020.

[50] Ian Goodfellow, Yoshua Bengio, and Aaron Courville. Deep learning. MIT press, 2016.

[51] Siavash Mirarab, Md Shamsuzzoha Bayzid, Bastien Boussau, and Tandy Warnow. Statistical binning enables an accurate coalescent-based estimation of the avian tree. Science, 346(6215):1250463, 2014.

[52] Md Shamsuzzoha Bayzid, Siavash Mirarab, Bastien Boussau, and Tandy Warnow. Weighted statistical binning: enabling statistically consistent genome-scale phylogenetic analyses. PLoS ONE, 10(6), 2015.

[53] Zhenxiang Xi, Liang Liu, Joshua S Rest, and Charles C Davis. Coalescent versus concatenation methods and the placement of amborella as sister to water lilies. Systematic Biology, 63(6):919–932, 2014.

[54] Chao Zhang, Maryam Rabiee, Erfan Sayyari, and Siavash Mirarab. Astral-iii: polynomial time species tree reconstruction from partially resolved gene trees. BMC Bioinformatics, 19(6):153, 2018.

[55] D.F. Robinson and L.R. Foulds. Comparison of phylogenetic trees. Mathematical Biosciences, 53:131–147, 1981.

[56] J H Degnan and N A Rosenberg. Gene tree discordance, phylogenetic inference and the multispecies coalescent. Trends in Ecology and Evolution, 26(6), 2009.

[57] Bent Fuglede and Flemming Topsoe. Jensen-shannon divergence and hilbert space embedding. In International Symposium on Information Theory, 2004. ISIT 2004. Proceedings., page 31. IEEE, 2004.

[58] Norman J Wickett, Siavash Mirarab, Nam Nguyen, Tandy Warnow, Eric Carpenter, Naim Matasci, Saravanaraj Ayyampalayam, Michael S Barker, J Gordon Burleigh, Matthew A Gitzendanner, et al. Phylotranscriptomic analysis of the origin and early diversification of land plants. Proceedings of the National Academy of Sciences, 111(45):E4859–E4868, 2014.

[59] Ning Zhang, Liping Zeng, Hongyan Shan, and Hong Ma. Highly conserved low-copy nuclear genes as effective markers for phylogenetic analyses in angiosperms. New Phytologist, 195(4):923–937, 2012.

[60] Siavash Mirarab and Tandy Warnow. Astral-ii: coalescent-based species tree estimation with many hundreds of taxa and thousands of genes. Bioinformatics, 31(12):i44–i52, 2015.

[61] Bryan T Drew, Brad R Ruhfel, Stephen A Smith, Michael J Moore, Barbara G Briggs, Matthew A Gitzendanner, Pamela S Soltis, and Douglas E Soltis. Another look at the root of the angiosperms reveals a familiar tale. Systematic Biology, 63(3):368–382, 2014.

[62] Vadim V Goremykin, Svetlana V Nikiforova, Patrick J Biggs, Bojian Zhong, Peter Delange, William Martin, Stefan Woetzel, Robin A Atherton, Patricia A Mclenachan, and Peter J Lockhart. The evolutionary root of flowering plants. Systematic Biology, 62(1):50–61, 2013.

[63] Erfan Sayyari and Siavash Mirarab. Fast coalescent-based computation of local branch support from quartet frequencies. Molecular Biology and Evolution, 33(7):1654–1668, 2016.

## References

[1] Md Shamsuzzoha Bayzid, Siavash Mirarab, Bastien Boussau, and Tandy Warnow. Weighted statistical binning: enabling statistically consistent genome-scale phylogenetic analyses. PLoS ONE, 10(6), 2015.

[2] Siavash Mirarab, Md Shamsuzzoha Bayzid, Bastien Boussau, and Tandy Warnow. Statistical binning enables an accurate coalescent-based estimation of the avian tree. Science, 346(6215):1250463, 2014.

[3] Erfan Sayyari and Siavash Mirarab. Fast coalescent-based computation of local branch support from quartet frequencies. Molecular Biology and Evolution, 33(7):1654–1668, 2016.

[4] Diederik P Kingma and Jimmy Ba. Adam: A method for stochastic optimization. arXiv preprint arXiv:1412.6980, 2014.

